# Distinguishing *PEX* gene variant severity for mild, severe, and atypical peroxisome biogenesis disorders in *Drosophila*

**DOI:** 10.1101/2024.11.14.623590

**Authors:** Vanessa A. Gomez, Oguz Kanca, Sharayu V. Jangam, Saurabh Srivastav, Jonathan C. Andrews, Michael F. Wangler

## Abstract

Peroxisomal biogenesis disorders (PBD) are autosomal recessive disorders caused by loss-of-function mutations of one of the *PEX* genes responsible for peroxisomal formation. Impaired peroxisome assembly causes severe multisystemic failure with patient phenotypes ranging from epilepsy, liver disease, feeding issues, biochemical abnormalities, and neurodegeneration. Variants in the same *PEX* gene can produce wide differences in severity, ranging from individuals with death in the first year of life to adults with milder complications. To study this strong genotype-phenotype correlation, we selected specific human *PEX* gene mutations and utilized *Drosophila* as a model organism. We generated flies replacing the coding sequence of our *Pex* gene of interest with a *KozakGAL4 (KZ)* promoter trap sequence. These cassettes simultaneously knock-out of the *Pex* gene and knock-in a *GAL4* driver, ideal for making “humanized” flies in which the human *PEX* gene can replace the fly loss. We assessed *Pex2^KZ^* and *Pex16^KZ^* lines in lifespan, bang sensitivity, and climbing assays and confirmed that these are strong loss-of-function alleles. In parallel, we generated human reference and variant UAS-cDNA lines of *PEX2* and *PEX16* variants in *Drosophila*. We observed nearly complete phenotypic rescue of *Drosophila Pex2* and *Pex16* loss when human *PEX2^Ref^*or *PEX16^Ref^*, respectively, were expressed. We also provide evidence for an allele severity spectrum in *PEX2* and *PEX16* in which some missense alleles, such as *PEX2^C247R^*, are equally severe as early truncations, such as *PEX2^R119*^*. We also observed that alleles associated with mild PBD, such as *PEX2^E55K^*, show variability depending on the assay but do not fully rescue. Finally, alleles associated with atypical ataxia phenotypes, such as *PEX16^F332Del^*, can perform as well as *PEX16^Ref^*, depending on the assay. Altogether, these *Drosophila* lines effectively model the range of severity of peroxisomal biogenesis disorders.

## Introduction

Peroxisomes are ubiquitous organelles that play an important role in cellular metabolism and perform specific biochemical functions in the cell, primarily involving complex lipids in eukaryotic cells^1,2^. Important biochemical functions performed by peroxisomes include fatty acid β-oxidation of very-long-chain fatty acids (VLCFA)^3^, α-oxidation of branched-chain fatty acids^4,5^, plasmalogen biosynthesis^6,7^, and catabolism of reactive oxygen species^8^ and glyoxylate^9,10^.

There are two main disorders associated with peroxisomes in humans: Peroxisomal biogenesis disorders (PBD) and single enzyme/protein deficiency disorders (SEPD)^11^. Peroxisomal biogenesis disorders are a group of autosomal recessive disorders caused by loss-of-function mutations in one of the peroxisomal matrix genes responsible for peroxisomal assembly and function^2,12^. Patients with PBD Zellweger spectrum disorders (ZSD) experience a wide range of multisystemic symptoms often within the first year of life, involving brain, bone, kidney, and liver, which can lead to death^13,14^. PBD consists of a spectrum of disorders with varying severity, ranging from mild to moderate to severe phenotypes^11^. Historically, severe PBD-ZSD (a.k.a. Zellweger syndrome) is the most dramatic and rapidly progressive disorder with a high mortality rate^11^. Intermediate PBD-ZSD (a.k.a. Neonatal adrenoleukodystrophy) has an infantile presentation with feeding problems and brain white matter changes, while mild PBD-ZSD (a.k.a. Infantile Refsum disease) is marked by hearing loss and retinal degeneration with a much milder cognitive impact^11,15^.

The peroxisomal biogenesis machinery is a conserved process, which is highly dependent on the action of 14 peroxins encoded by *PEX* genes that are required for matrix protein import, peroxisome membrane assembly, and peroxisome proliferation^16,17^. Early peroxisomal proteins including *PEX3*, *PEX19*, and *PEX16* aid in the designation of an ER-derived lipid bilayer and its maturation to a pre-peroxisomal vesicle^17^. Membrane proteins are then incorporated to allow enzyme import into the maturing peroxisome with the help of *PEX2* and the importer complex^18^. Mature peroxisomes can then perform a wide range of biochemical functions important for cell function and proper metabolism^1,2^. Therefore, it is critical to study *PEX* genes involved in different aspects of biogenesis, such as *PEX2* and *PEX16*, when characterizing peroxisomal biogenesis disorders.

The severity spectrum of PBD-ZSD correlates to the allele severity of the *PEX* gene mutations. In general, clinical severity correlates to lower levels of residual *PEX* gene activity and much less to which *PEX* gene is involved (e.g. *PEX16* vs. *PEX2* vs. *PEX3*)^19,20^. Patients with biallelic loss of function ‘null’ *PEX* alleles are more likely to exhibit drastic biochemical defects and severe cases of PBD-ZSD than patients harboring one of the hypomorphic alleles that retain partial PEX protein functions. For instance, milder alleles of *PEX2* such as *PEX2^E55K^*in compound heterozygotes *(PEX2^Null^/PEX2^E55K^)* are associated with mild PBD-ZSD, where the phenotypes are mild or intermediate due to residual *PEX2* function^21,22^. Likewise, we also observed a case with a single amino acid deletion of *PEX16* with normal plasma VLCFA and an atypical ataxia phenotype^23,24^. In summary, PBD-ZSD exhibits a clinical spectrum that correlates with specific alleles.

Peroxisomal biology is highly conserved across eukaryotes, and recent studies have demonstrated the evolutionary conservation of peroxisomal biogenesis in *Drosophila*^25–30^. Previous studies have found that *Pex* mutations in *Drosophila* alter lipid metabolism, muscle function, and spermatogenesis^25–27^. Our research has focused on *Pex2* and *Pex16* mutant flies, which has allowed us to compare different biogenesis defects. We previously documented a detailed phenotypic characterization in flies on two alleles for both *Pex2* and *Pex16*, studying the mutants in trans with genomic deficiencies, and creating genomic rescue strains for each mutation^30^. We also showed that the peroxisomes were similarly functionally and morphologically defective in *Pex2* or *Pex16* mutants, and mutant flies had short lifespans, increased bang sensitivity, lacked flight ability, and showed reduced activity^30^. Additionally, functional analysis of peroxisomal lipids allowed for a comprehensive study of VLCFA metabolites in different stages of development, as well as findings of a dramatic loss of plasmalogen synthesis in both *Pex2* and *Pex16* mutant flies^30^. Altogether, previous studies of peroxisomes in *Drosophila* support the high conservation of the peroxisomal biogenesis machinery in flies and humans.

A key question that has not been addressed previously is whether human transgenes could rescue deleterious *Pex* mutant phenotypes in flies. While comprehensive studies have functionally characterized the conservation of the peroxisomal biogenesis machinery in flies and phenotypes of *Pex* mutants, functional studies on known human proband variants involved in PBD in the context of the human protein have not yet been done. Here, we utilize *KozakGAL4* cassettes to knock-out *Pex* genes of interest, creating an effective null background that allowed us to express human reference and variant UAS-cDNAs of the targeted gene to assess the rescue of the *Pex* mutants. We assayed the flies for phenotypes that allowed us to study disease progression resulting from the expression of the human variants that could be classified as pathogenic or likely pathogenic. We also studied whether the human protein can functionally replace the loss of the endogenous fly Pex protein. With these findings, we have found a method to functionally characterize individual human *PEX* mutations in *Drosophila* and create an allelic series.

## Results

### Generation of the *KozakGAL4* lines resulting in a *GAL4* gene trap allele and examination of the spatiotemporal expression pattern

To study the neurodevelopmental and neurodegenerative phenotypes seen in PBD patients, we generated novel alleles for our genes of interest with *KozakGAL4* cassettes in *Drosophila.* This strategy replaces the coding sequence of genes with the *KozakGAL4* cassette using CRISPR induced homologous recombination using sgRNAs targeting the 5’ and 3’ UTR^31^. These alleles are useful for determining the gene’s expression pattern, studying the effect of loss-of-function of the gene product, assessing protein subcellular localization with the help of a GFP protein trap, and expression of UAS-cDNAs of the targeted gene and variants to assess rescue of the mutant phenotypes^31^.

We have generated fly lines in which the coding sequence of *Pex2* and *Pex16* respectively were replaced by the *KozakGAL4* gene trap sequence (**Figure 1A-B**). We confirmed the insertions by PCR. This technology leads to a null allele (*Pex2^KZ^* and *Pex16^KZ^*) with expression of *GAL4* in a similar spatial and temporal pattern as the protein encoded by the targeted gene^31^. This method has been successfully employed to study expression patterns and to “humanize” the gene by driving the human cDNA to effectively replace the fly loss^32,33^.

**Figure 1.**
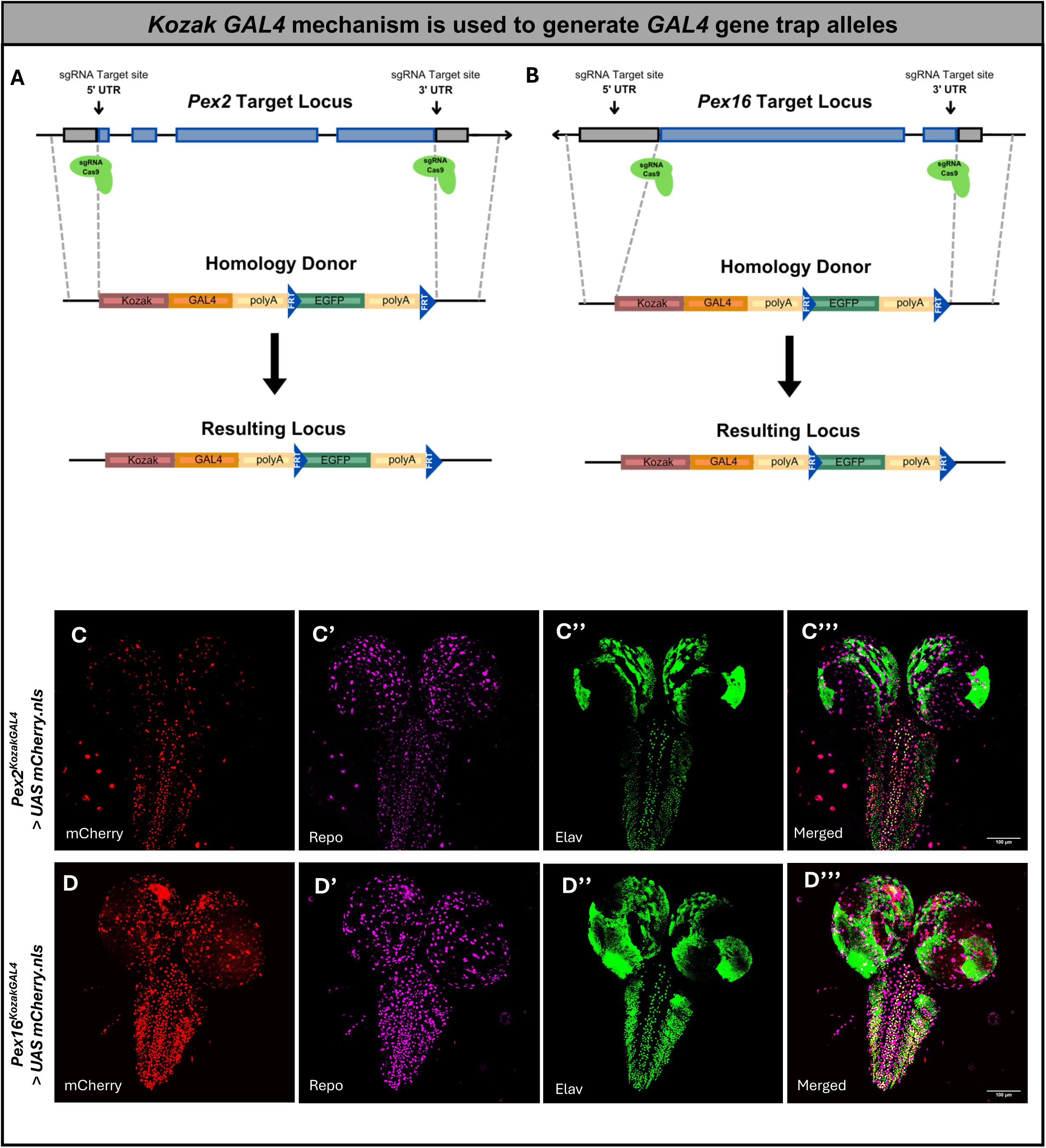
The *KozakGAL4* knock-in/knock-out strategy, which replaces the coding region of the *Pex* gene of interest by identifying sgRNA target sites in the 5’ UTR and 3’ UTR. Gray boxes, UTRs; blue boxes, *Pex*-coding region. (A) Schematics of *Pex2* locus and targeting strategy. (B) Schematics of *Pex16* locus and targeting strategy. (C) Expression pattern of *Pex2^KZ^* in 3rd instar larval brain. (C’) Repo expression. (C’’) Elav expression. (C’’’) Merged image with co-localization within the ventral nerve cord, indicating *Pex2^KZ^* expression in both neurons (Elav) and nuclear glia (Elav) in the larval brain. (D) Expression of *Pex16^KZ^* expression pattern in the third instar *Drosophila* larval brain. (D’) Repo expression. (D’’) Elav expression. (D’’’) Merged image with co-localization within the ventral nerve cord, indicating *Pex16^KZ^* expression in both neurons (Elav) and nuclear glia (Elav) in the larval brain.

We examined the expression using third instar larval *Drosophila* brain (**Figure 1C-D**) and assessed whether the *Pex2* and *Pex16* genes were expressed in neurons, glia, or both by co-staining with Repo and Elav. Within the ventral nerve cord (VNC), we observed co-staining of cells expressing *Pex2* or *Pex16* with both Repo and Elav (**Figure 1C-D**). We noted that *Pex2* expression was generally sparser in the larval third instar brain. Altogether, *Pex2* and *Pex16* appear to be expressed in both neurons and glia in the larval brain. The novel alleles *Pex2^KZ^* and *Pex16^KZ^* behave similar to known *Pex2* and *Pex16* null

### alleles

Having generated these unique *KozakGAL4* cassettes, we wanted to study the strength of these loss-of-function alleles of *Pex2* and *Pex16* using behavior assays. We crossed *Pex2^KZ^* to known *Pex2* null alleles, including a transposable element, a deletion allele, and 2 frameshift alleles *Pex2*^1^ and *Pex2*^2^ (**Figure 2A**). Similarly, we crossed *Pex16^KZ^* to a known *Pex16* null allele, *Pex16*^1^ (**Figure S1A**).

**Figure 2.**
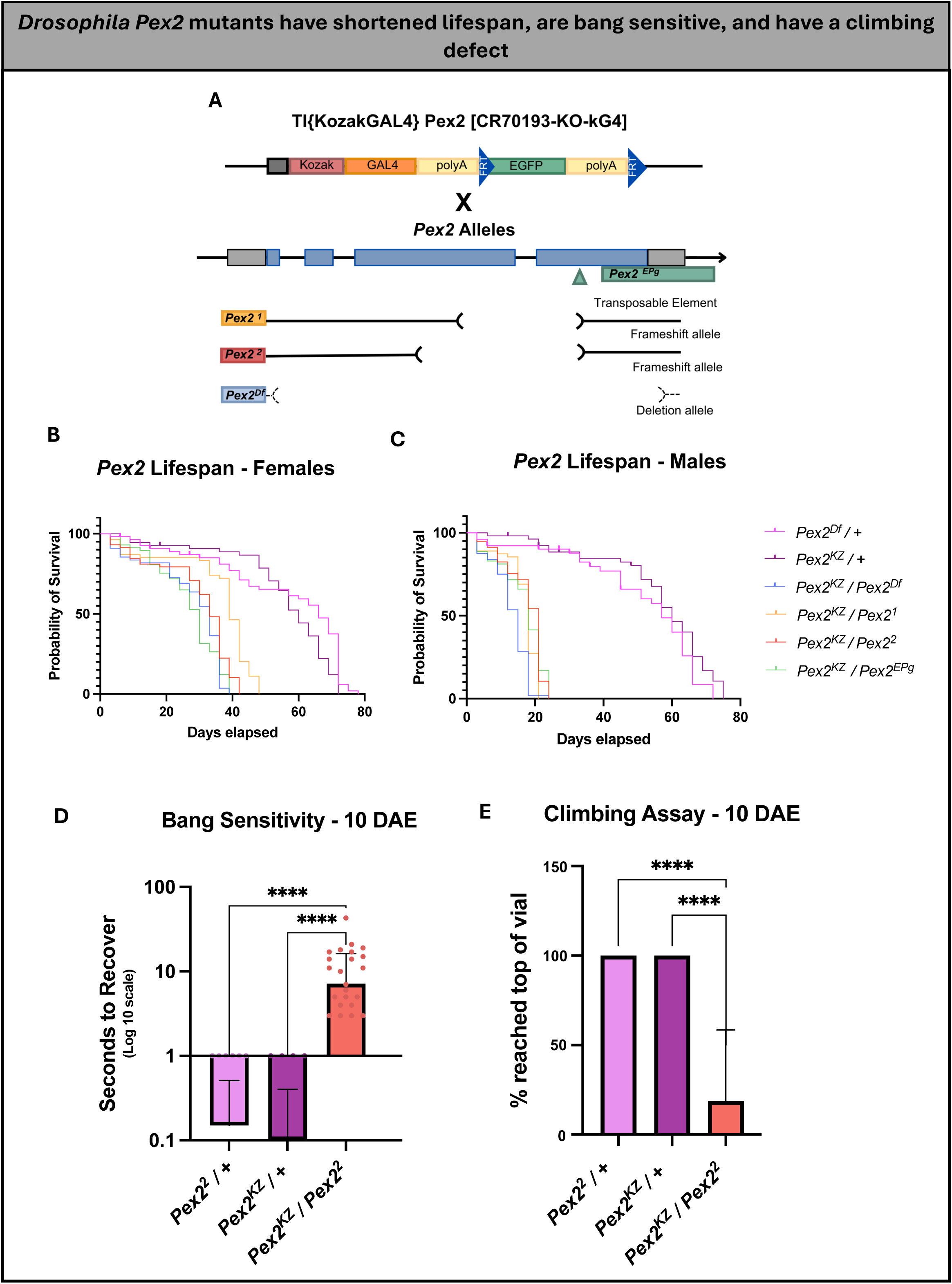
*Drosophila Pex2* mutants have shortened lifespan, are bang-sensitive, and have a climbing defect. (A) Schematic representation of fly *Pex2* gene along with four alleles, including a coding P-element insertion (*Pex2^EPg^*), 2 deletion alleles (*Pex2^1^*, *Pex2^2^*), and *Pex2*-*KozakGAL4 (Pex2^KZ^*). (B) *Pex2* female lifespan assay shows that the *Pex2* mutants have a shorter lifespan compared to control lines (pink and purple). (C) *Pex2* male lifespan assay shows that the *Pex2* mutants have a shorter lifespan compared to control lines, worse than female mutant flies. (D) *Pex2* null flies have a significant bang-sensitive phenotype (red) compared to controls (pink and purple) observed at 10 days after eclosion (DAE). (E) *Pex2* null flies have a significant climbing deficiency (red) compared to controls (pink and purple) observed at 10 days after eclosion. [* = *p*-value is less than 0.05. ** = *p*-value is less than 0.01. *** = *p*-value is less than 0.001. **** = *p*-value is less than 0.0001]

We first analyzed the lifespan of the F1 progeny of the flies crossed from *Pex2^KZ^* to the 4 known *Pex2* null alleles. Lifespan analysis showed a significant difference between experimental flies and controls in females (**Figure 2B**). While both control genotypes had an average lifespan of 55.5 days, female flies *Pex2^KZ^/Pex2^Df^* lived an average of 27.2 days, *Pex2^KZ^/Pex2*^1^ lived 35.7 days, *Pex2^KZ^/Pex2*^2^ lived 29.5 days, and *Pex2^KZ^/Pex2^EPg^*lived an average of 26.6 days. This effect was even more dramatic in males (**Figure 2C**). Control male flies *Pex2^Df^* /+ and *Pex2^KZ^* /+ lived an average of 49.3 days and 55.9 days, respectively. Meanwhile, male flies *Pex2^KZ^/Pex2^Df^* lived an average of 12.8 days, *Pex2^KZ^/Pex2^1^* lived 16.4 days, *Pex2^KZ^/Pex2^2^*lived 17.4 days, and *Pex2^KZ^/Pex2^EPg^* lived an average of 16.5 days.

We also assessed bang sensitivity and climbing, focusing on the *Pex2^KZ^/Pex2^2^*null flies compared to controls. Bang sensitivity is an assay used to test for seizure-like behavior and paralysis following mechanical stimulation^34^. 10 days after eclosion (DAE), the *Pex2^KZ^/Pex2^2^*null flies were found to have a significant bang-sensitive phenotype compared to heterozygous controls (**Figure 2D**). The climbing assay evaluates the flies’ natural tendency to climb, which is known as negative geotaxis^35^. In *Drosophila*, negative geotaxis relies on the presence of intact sensory and motor systems, and a defect in climbing may be an effective readout for peroxisomal disease progression in the fly^35^. When observing the climbing behavior of the *Pex2^KZ^/Pex2^2^* flies compared to heterozygous controls at 10 days after eclosion, we observed only 18% of *Pex2* null flies had the ability to climb a full 8 cm within 60 seconds, demonstrating a clear lack of climbing ability and putative disease progression (**Figure 2E**).

In parallel, we performed lifespan analysis on the F1 progeny of the flies crossed from *Pex16^KZ^* to the *Pex16^1^* null allele. Lifespan analysis showed a significant difference between the *Pex16^KZ^/Pex16^1^* and controls in both females and males (**Figure S1B-C**). Control female flies *Pex16^1^*/+ and *Pex16^KZ^/+* lived an average of 49 and 50.5 days, respectively. Meanwhile, *Pex16^KZ^/Pex16^1^* null female flies lived an average of 13.2 days. Control male flies *Pex16^1^*/+ and *Pex16^KZ^/+* lived an average of 40 and 51.2 days, respectively. Meanwhile, *Pex16^KZ^/Pex16^1^*null male flies lived an average of 8.5 days.

When performing the bang sensitivity assay on *Pex16^KZ^/Pex16^1^*null flies, we found a significant seizure-like phenotype that was not seen in the heterozygous control flies (**Figure S1D**). When assessing climbing ability 10 days after eclosion, only 3% of the *Pex16^KZ^/Pex16^1^* flies were able to climb the full 8 cm within 60 seconds (**Figure S1E**). In summary, the *KozakGAL4* cassette alleles *Pex2^KZ^* and *Pex16^KZ^* behave as genetic null alleles similar to previously documented deletions of *Pex2^2^* and *Pex16^1^* in flies.

### Rescue-based humanization of *Drosophila Pex* genes to study rare *PEX* variants

Having established that the *Pex2^KZ^* and *Pex16^KZ^* are strong loss-of-function alleles and express transgene under *GAL4*, we chose to further study rare human variants involved in PBD. The rescue-based humanization experiments of *Pex2* and *Pex16* involve the generation of transgenic *Drosophila* lines containing human reference and variant *PEX2* and *PEX16* under the control of the UAS^36,37^. “Humanization” was performed by expressing the reference or variant UAS-human cDNA in either the *Pex2^KZ^/Pex2^2^* or *Pex16^KZ^/Pex16^1^*mutant background, as appropriate, and observing for differences in phenotype that could suggest a *Pex*-specific effect^36,37^.

*PEX2* is found on chromosome 8 (**Figure 3A**) and contains 5 transmembrane domains and a Zinc finger binding domain (**Figure 3B**). Autosomal recessive mutations in *PEX2* have been associated with a range of mild to intermediate to severe cases of PBD-ZSD. We selected four variants that span the range of clinical severity to test in *Drosophila*^21,22,38–42^. We generated transgenic *Drosophila* lines with the four selected human variants, as well as a human reference line, designed to express the human protein (codon optimized) in *Drosophila* (**Figure 3C**). Our first variant, *PEX2^E55K^*, was classified on ClinVar (Variation ID:13705) as a likely pathogenic variant in an individual with mild PBD-ZSD who was compound heterozygous with a pathogenic missense variant in *PEX2^R1^*^19^*\** ^21,22^. Experimental evidence from cells from patients with this particular *PEX2* variant demonstrates that the *PEX2^E55K^* variant displays residual enzymatic activity and is competent to import target proteins into peroxisomes^22,38^. The *PEX2^E55K^* cells also demonstrate “peroxisomal mosaicism” in cell culture, where some cells within the culture do not contain peroxisomes^22,38^. This feature is thought to be related to the mild temperature sensitivity of the *PEX* allele and presumed stochastic factors within the cell culture^22,38^. *PEX2^R119*^* is noted on ClinVar (Variation ID: 13704) as a pathogenic variant in several PBD-ZSD cases and is known to alter peroxisome assembly^39^. When homozygous, the *PEX2^R119*^*variant has been shown to cause a severe PBD-ZSD (Zellweger syndrome) phenotype and death in early infancy^40^. When seen in cases with other PBD-ZSD variants, the clinical severity of the *PEX2^R119*^* variant varies based on the other allele, with findings of mild PBD-ZSD with *PEX2^R119*^/ PEX2^E55K^* and findings of mild PBD-ZSD manifesting as childhood-onset cerebellar ataxia and an axonal sensorimotor polyneuropathy with *PEX2^R119*^* and a 1-bp insertion (c.865_866insA;170993.0006) in the *PEX2* gene^22,41^. The *PEX2^W223*^* variant is also noted on ClinVar (Variation ID: 139590) as a pathogenic variant. The proband, who was homozygous for the *PEX2^W223*^* variant, displayed mild PBD-ZSD and had normal infancy, but by the age of 22 months, he had hypotonia, cerebellar and vermian atrophy, and continued to deteriorate until he died at age 13^40,42^. The last *PEX2* variant generated was *PEX2^C247R^* mutation, which is also noted on ClinVar (Variation ID: 139589) as a pathogenic variant. This variant was seen in a newborn patient with severe PBD-ZSD (Zellweger syndrome), with a low birth weight, severe hypotonia, seizures, absent corpus callosum, severe icterus, and absent peroxisomes leading to death at 3 months of age^40^.

**Figure 3.**
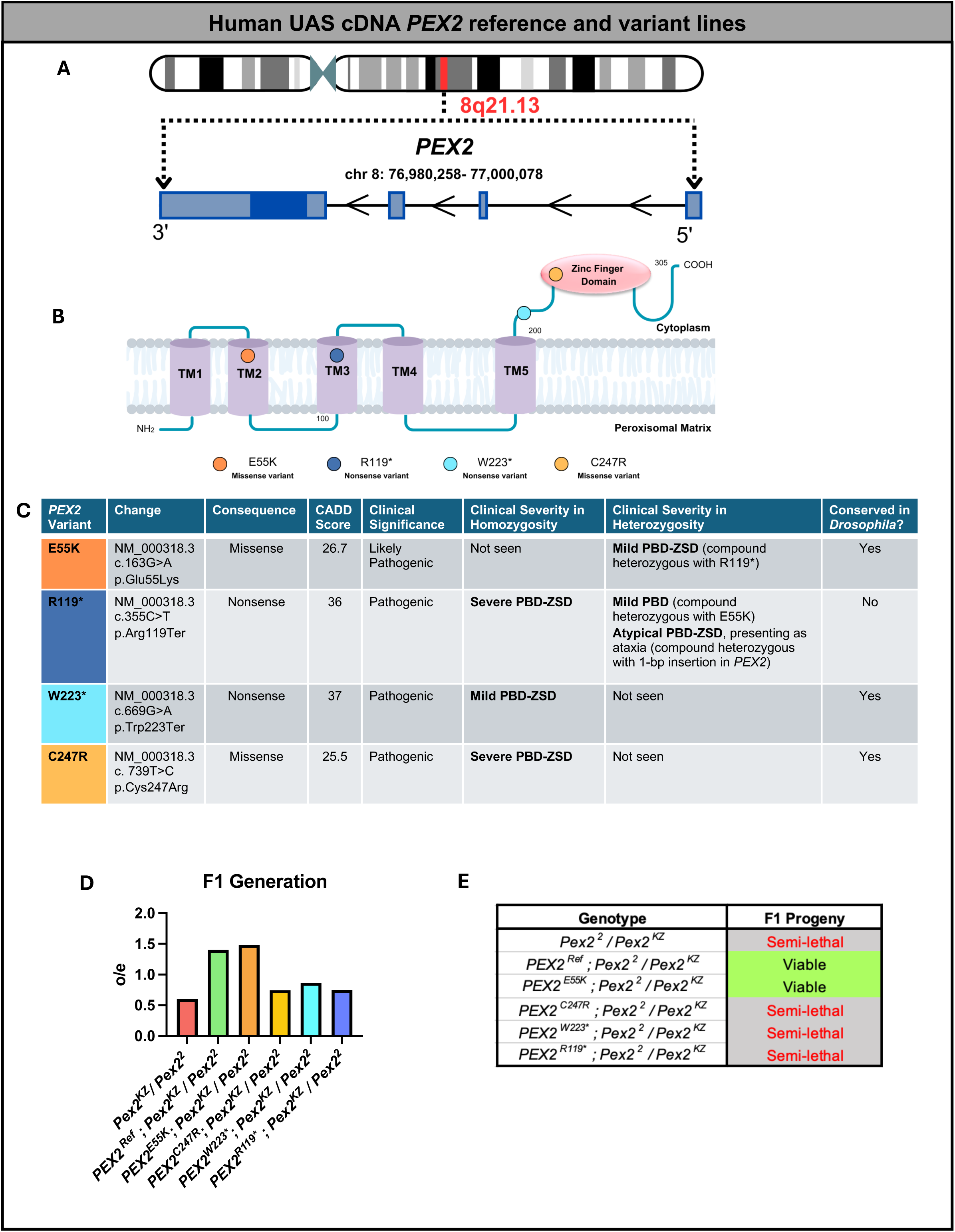
Human UAS cDNA *PEX2* reference and variant lines. (A) Schematic representation of human *PEX2* gene. (B) Schematic representation of human PEX2 protein and variant locations. (C) *PEX2* variant table indicates the consequence of the change, pathogenicity prediction, clinical significance, clinical severity in homozygosity and heterozygosity, and conservation in *Drosophila.* (D) Indicates the observed/expected Mendelian ratio of the F1 generation of human *PEX2* variants, in a fly null background. (E) Assessment of the phenotype of indicated genotypes as lethal, viable, or semi-lethal.

Once we generated the reference and variant lines, we then crossed these lines to the *Pex2^2^* null background and generated flies that were compound heterozygous *Pex2^KZ^/Pex2^2^*expressing the human transgenes. We had previously observed that *Pex2^KZ^/Pex2^2^*without transgenes were semi-lethal leading to approximately 50% of the expected progeny by Mendelian ratios (**Figure 3D**). The *PEX2^Ref^* human transgene was able to fully rescue this semi-lethality. Interestingly, the *PEX2^E55K^* also fully rescued but the *PEX2^C247R^*, *PEX2^W223*^* and *PEX2^R119*^* did not appear to rescue the semi-lethality, although *PEX2^W223*^* was above 80% observed/expected (**Figure 3D-E**).

We also followed a similar strategy to study human variants for *PEX16* **(Figure S2)**. *PEX16* is found on chromosome 11 (**Figure S2A**) and contains 2 transmembrane domains, a peroxisomal location domain, and a *PEX19* interaction domain (**Figure S2B**). We have generated 2 transgenic *Drosophila* lines with human variants of varying clinical severity, as well as a *PEX16* human reference line (**Figure S2C**). Our variant *PEX16^R176*^* is interpreted as a pathogenic variant on ClinVar (Variation ID: 6466) and is expected to result in an absent or disrupted protein. This variant has been observed as homozygous or in trans with another pathogenic variant in patients with the hallmark clinical features of Zellweger syndrome^43,44^. We also generated a transgene for a unique case that we have previously reported. The patient was homozygous for a *PEX16^F332del^* variant that is currently interpreted as likely pathogenic on ClinVar (Variation ID: 209181)^23,24,45^. Our proband had no symptoms until the age of 7 and then displayed a progressive ataxia phenotype that was undiagnosed for 18 years as the patient had multiple plasma studies including peroxisomal biochemical assays that had normal findings^23,24^.

Utilizing both variants along with the reference line in *PEX16* we documented rescue of semi-lethality of *Pex16^KZ^/Pex16^1^*, and complete rescue by *PEX16^Ref^*(**Figure S2D**). For the two *PEX16* alleles, we saw no rescue of semi-lethality for the *PEX16^R176*^* but the *PEX16^F332del^*fully rescued (**Figure S2D-E**).

### Human *PEX* reference gene rescues fly peroxisome morphology

Next, we wanted to test the impact of the human proteins on rescue to peroxisomal morphology defects (**Figure 4**). We utilized anti-Pex3 staining in the 3^rd^ instar larval body wall muscle. The Pex3 antibody stains the early pre-peroxisomal vesicles, and we have previously observed that, in *Pex2* null and *Pex16* null mutants, Pex3 puncta are present but are significantly reduced with a more severe reduction in *Pex16* mutants^25,30^.

**Figure 4.**
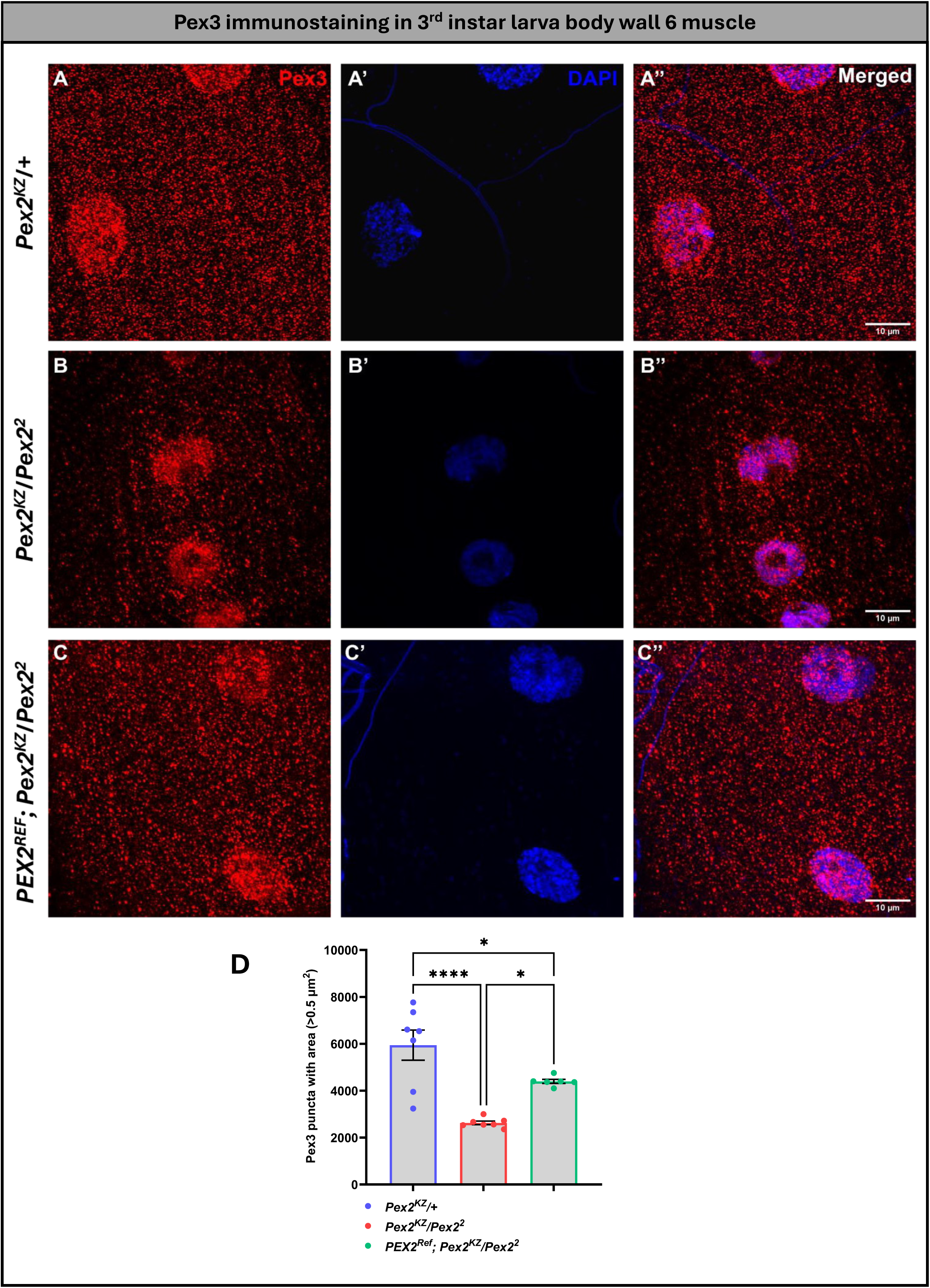
Human *PEX2* expression in *Pex2* null background larvae significantly rescues peroxisomes in 3^rd^ instar larva body wall 6 muscle. (A, A’& A’’) represents control group; (B, B’, & B’’) represents *Pex2* null flies & (C, C’ & C’’) represents human rescue group. Quantification of Pex3 positive puncta between the three genotypes. [*p<0.05; ****p<0.0001]

In the *Pex2* control line, we see Pex3 staining in a puncta pattern throughout the body wall (**Figure 4A**). For *Pex2^KZ^/Pex2^2^* mutants, we observed a dramatic reduction in the Pex3 puncta compared to control (*Pex2^KZ^/+*) (**Figure 4B, 4D**). This peroxisomal morphology phenotype is partially rescued by the human *PEX2* expression (**Figure 4C**). The Pex3 reduction in the *Pex2^KZ^/Pex2^2^*mutant is dramatic and statistically significant (*p*<0.0001) providing further evidence that the *Pex2^KZ^* is a strong loss-of-function allele (**Figure 4D**). The human reference *PEX2* transgene expression leads to a robust increase in the Pex3 puncta compared to the *Pex2^KZ^/Pex2^2^* mutant, but it is also significantly less than the control indicating a partial rescue (**Figure 4D**). In the *Pex16* lines, we also saw a dramatic loss of Pex3 puncta in the mutant compared to the control, and this phenotype was also partially rescued by the human *PEX16* transgene (**Figure S3**).

In summary, both human *PEX2* and *PEX16* human genes can function in *Drosophila* and partially rescue Pex3 staining, indicating that the human genes can successfully restore peroxisomes in a fly mutant. Therefore, we have generated humanized flies for *PEX2* and *PEX16* in which disease-causing variants and other phenotypes can be tested.

### Human *PEX* reference partially rescues fly *Pex* null behavior phenotypes, but the variants fail to do so

To further study the effects of rescue-based humanization of *Pex* genes, we moved to three behavior assays (lifespan, bang sensitivity, and climbing) to observe and compare the behavior of the *PEX* variants and reference lines. We performed a lifespan analysis of our humanized *PEX2* variants and reference flies (**FIGURE 5A-B**). We noted that the human *PEX2^Ref^* partially rescues the shortened lifespan of *Pex2^KZ^/Pex2^2^* in females and there is some degree of rescue with the *PEX2^E55K^* (**FIGURE 5A**). The severe loss-of-function variants (*PEX2^R119*^, PEX2^C247R^, PEX2^W223*^)* have a similar lifespan to the *Pex2^KZ^/Pex2^2^*mutant and fail to rescue (**FIGURE 5A**). Female control flies *Pex2^KZ^/+* lived an average of 55.5 days and *Pex2^2^/+* lived an average of 40.7 days. Female mutant flies *Pex2^KZ^/Pex2^2^*lived an average of 29 days, *PEX2^Ref^; Pex2^KZ^/Pex2^2^*lived an average of 33.1 days, *PEX2^E55K^;Pex2^KZ^/Pex2^2^*lived an average of 25.9 days, *PEX2^C247R^;Pex2^KZ^/Pex2^2^*lived an average of 17.2 days, *PEX2^W223*^;Pex2^KZ^/Pex2^2^*lived an average of 21.2 days, and *PEX2^R119*^;Pex2^KZ^/Pex2^2^*lived an average of 19.3 days. In males, the lifespan for *Pex2^KZ^/Pex2^2^*is even shorter and is not rescued by the severe loss-of-function variants, but the *PEX2^Ref^* appears to rescue. Interestingly, the mild allele *PEX2^E55K^* is indistinguishable from null in the male lifespan assay (**FIGURE 5B**).

**Figure 5.**
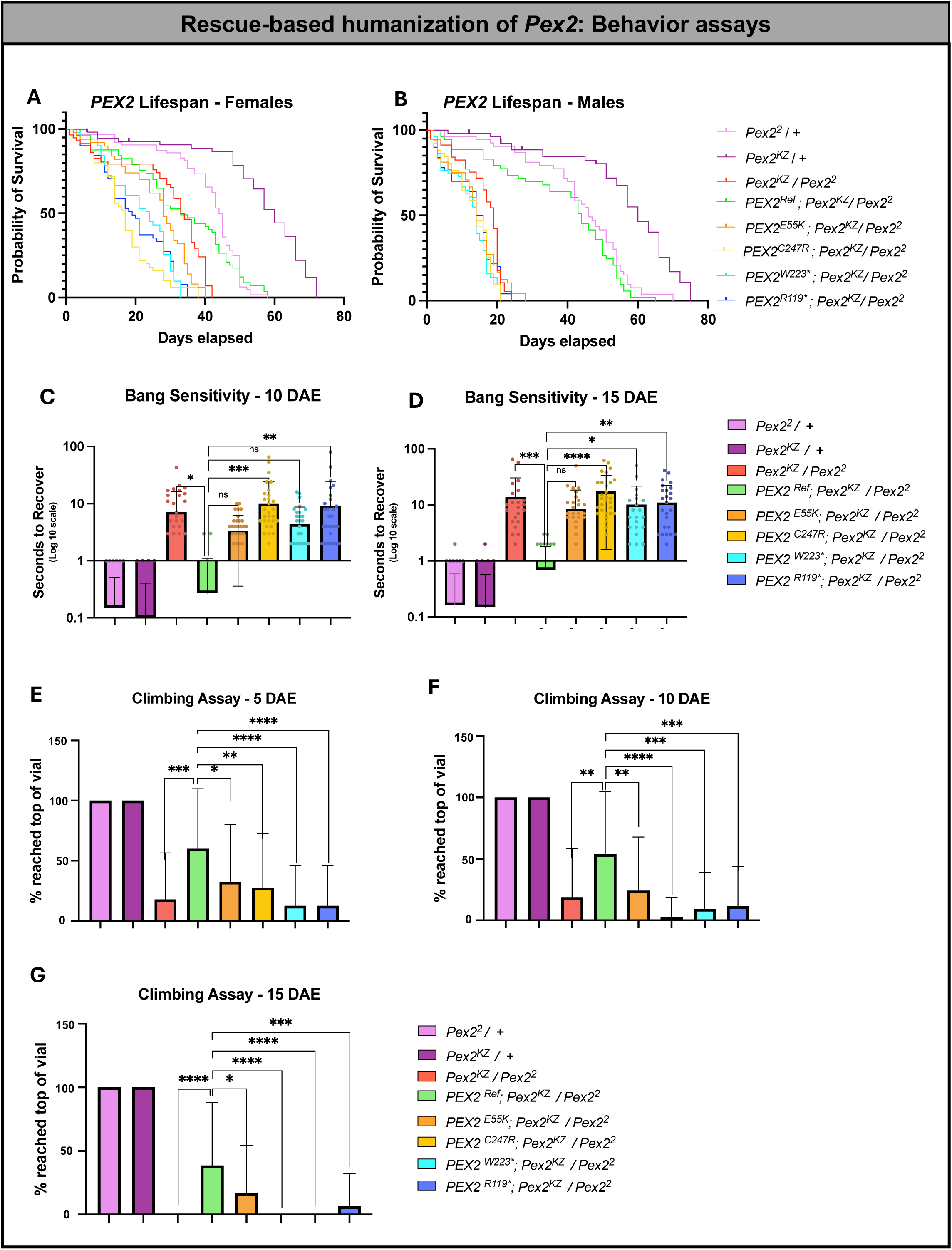
Rescue-based humanization of *Pex2*: Behavior Assays. (A) Lifespan analysis of *PEX2^Ref^ ^and^ ^Variant^* female flies, along with our *Pex2* null and control lines. (B) Lifespan analysis of *PEX2^Ref^ ^and^ ^Variant^* male flies, along with our *Pex2* null and control lines. (C) Bang sensitivity assay of *PEX2^Ref^ ^and^ ^Variant^* female flies, along with our *Pex2* null and control lines, at 10 days after eclosion (DAE). (D) Bang sensitivity assay of *PEX2^Ref^ ^and^ ^Variant^* female flies, along with our *Pex2* null and control lines, at 15 days after eclosion. (E-G) Climbing assay of *PEX2^Ref^ ^and^ ^Variant^*female flies, along with our *Pex2* null and control lines, at 5 days, 10 days, and 15 days after eclosion. [* = *p*-value is less than 0.05. ** = *p*-value is less than 0.01. *** = *p*-value is less than 0.001. **** = *p*-value is less than 0.0001]

Additionally, the male reference flies have a much more significant rescue, especially noting that *Pex2* male null flies tend to have a shorter lifespan than female nulls. Male control flies *Pex2^KZ^/+* lived an average of 55.9 days and *Pex2^2^/+* lived an average of 43.9 days. Male mutant flies *Pex2^KZ^/Pex2^2^* lived an average of 16.2 days, *PEX2^Ref^; Pex2^KZ^/Pex2^2^*lived an average of 38.4 days, *PEX2^E55K^;Pex2^KZ^/Pex2^2^*lived an average of 13.8 days, *PEX2^C247R^;Pex2^KZ^/Pex2^2^*lived an average of 13 days, *PEX2^W223*^;Pex2^KZ^/Pex2^2^*lived an average of 12.4 days, and *PEX2^R119*^;Pex2^KZ^/Pex2^2^*lived an average of 13.2 days. Next, we observed the *PEX2* variants and reference flies’ behavior in response to the bang sensitivity assay at 10 and 15 days after eclosion (**FIGURE 5C-D**). On day 10, we found that *PEX2^Ref^* can partially rescue the *Pex2* null phenotype. However, the variants *PEX2^R119*^* and *PEX2^C247R^* fail to rescue, and *PEX2^E55K^*and *PEX2^W223*^* did not differ significantly from the reference (**FIGURE 5C**). At day 15, the flies become slightly more bang-sensitive, showing that there is progressive neuronal dysfunction (**FIGURE 5D**). Interestingly, *PEX2^E55K^*continued to behave not significantly different to *PEX2^Ref^* at day 15 (**FIGURE 5D**). We also noted age-dependent worsening of phenotype when observing the flies’ ability to climb at 5 days, 10 days, and 15 days after eclosion (**FIGURE 5E-G**). We found, again, that the *PEX2^Ref^*flies were able to partially rescue the fly null phenotype, and we found that the variants behave similarly to the *Pex2* null flies. *PEX2^R119*^, PEX2^C247R^*, and *PEX2^W223*^*expressed in flies all behave as null alleles in each of the three assays. However, we show that the *PEX2^E55K^* variant, associated with mild PBD-ZSD rescues lifespan for females but not males, rescues bang sensitivity, and has an intermediate rescue of climbing at different ages. Altogether, the *PEX2^E55K^* allele is likely a hypomorphic allele.

We conducted the same three behavior assays for the *PEX16* variants and reference lines. We first performed a lifespan analysis of our humanized *PEX16* variants and reference flies (**FIGURE S4A-B**). We noted that the human *PEX16^Ref^* partially rescues the shortened lifespan of *Pex16^KZ^/Pex16^1^* in both females and males, and there is also partial rescue with *PEX16^F332del^* (**FIGURE S4A-B**). The severe loss-of-function *PEX16^R176*^* variant has a similar lifespan to *Pex16^KZ^/Pex16^1^* and fails to rescue in both females and males (**FIGURE S4A-B**). Female control flies *Pex16^KZ^/+* lived an average of 50.5 days and *Pex16^1^/+* lived an average of 49 days. Female mutant flies *Pex16^KZ^/Pex16^1^*lived an average of 13.2 days, *PEX16^Ref^; Pex16^KZ^/Pex16^1^*lived an average of 26.2 days, *PEX16^F332del^;Pex16^KZ^/Pex16^1^*lived an average of 26.3 days, and *PEX16^R176*^;Pex16^KZ^/Pex16^1^*lived an average of 11 days. Male control flies *Pex16^KZ^/+* lived an average of 51.2 days and *Pex16^1^/+* lived an average of 40 days. Male mutant flies *Pex16^KZ^/Pex16^1^* lived an average of 8.5 days, *PEX16^Ref^; Pex16^KZ^/Pex16^1^*lived an average of 23 days, *PEX16^F332del^;Pex16^KZ^/Pex16^1^*lived an average of 26.2 days, and *PEX16^R176*^;Pex16^KZ^/Pex16^1^*lived an average of 12 days. Next, we performed the bang sensitivity assay on our *PEX16* variant and reference flies at 10 and 15 days after eclosion (**FIGURE S4C-D**). We found that our human reference flies could rescue the *Pex16* null bang sensitivity. Interestingly, the *PEX16^R176*^*variant flies could not rescue the bang-sensitive phenotype, but the *PEX16^F332del^*variant flies had no significant difference with the reference flies at day 10 or day 15 and could also rescue the phenotype. We also observed *PEX16^Ref^*and variant flies’ ability to climb at 5 days, 10 days, and 15 days (**FIGURE S4E-G**). We found that the *PEX16^R176*^* variant behaved similarly to the *Pex16* null flies and could not rescue the phenotype. However, the reference and *PEX16^F332del^* flies could partially rescue the climbing defect. Interestingly, the *PEX16^F332del^*flies were similar to *PEX16^Ref^* at 5 and 10 days and actually behaved better than *PEX16^Ref^* at day 15 (**Figure S4G**). In summary, the *PEX16^F332del^* allele which is associated with an atypical ataxia clinical presentation exhibits a complete rescue of shortened lifespan, a complete rescue of bang sensitivity at day 10 and day 15, a near complete rescue of bang sensitivity and Day 15, and overall better rescue than reference for climbing.

### Loss of *Pex* genes leads to phenotypes in the direct flight muscle, but the human *PEX* reference is able to rescue

In our previous characterization of *Pex2* and *Pex16* deletion alleles, we observed climbing, lifespan, and flight defects^30^. Given that we saw locomotor and lifespan rescue with the human transgenes, we decided to assess the morphology of indirect flight muscles. In control flies (*Pex2^KZ^/+*), we observe Pex3 puncta intercalating the muscle fibers and stained HRP to examine the innervating nerve fiber in 0-day-old flies (**Figure 6A**). In the mutant, we observed a reduction in Pex3 staining and an apparent thickening of the surrounding nerve fiber, an intriguing observation as the 0-day-old flies exhibit locomotor defects in our bang sensitivity and climbing assay (**Figure 6B**). The human *PEX2* transgene expression partially rescued peroxisomes (Pex3 staining) and appears to rescue the nerve fiber thickening observed in the mutant (**Figure 6C**). The differences were statistically significant, indicating a partial peroxisome rescue (**Figure 6D**).

**Figure 6.**
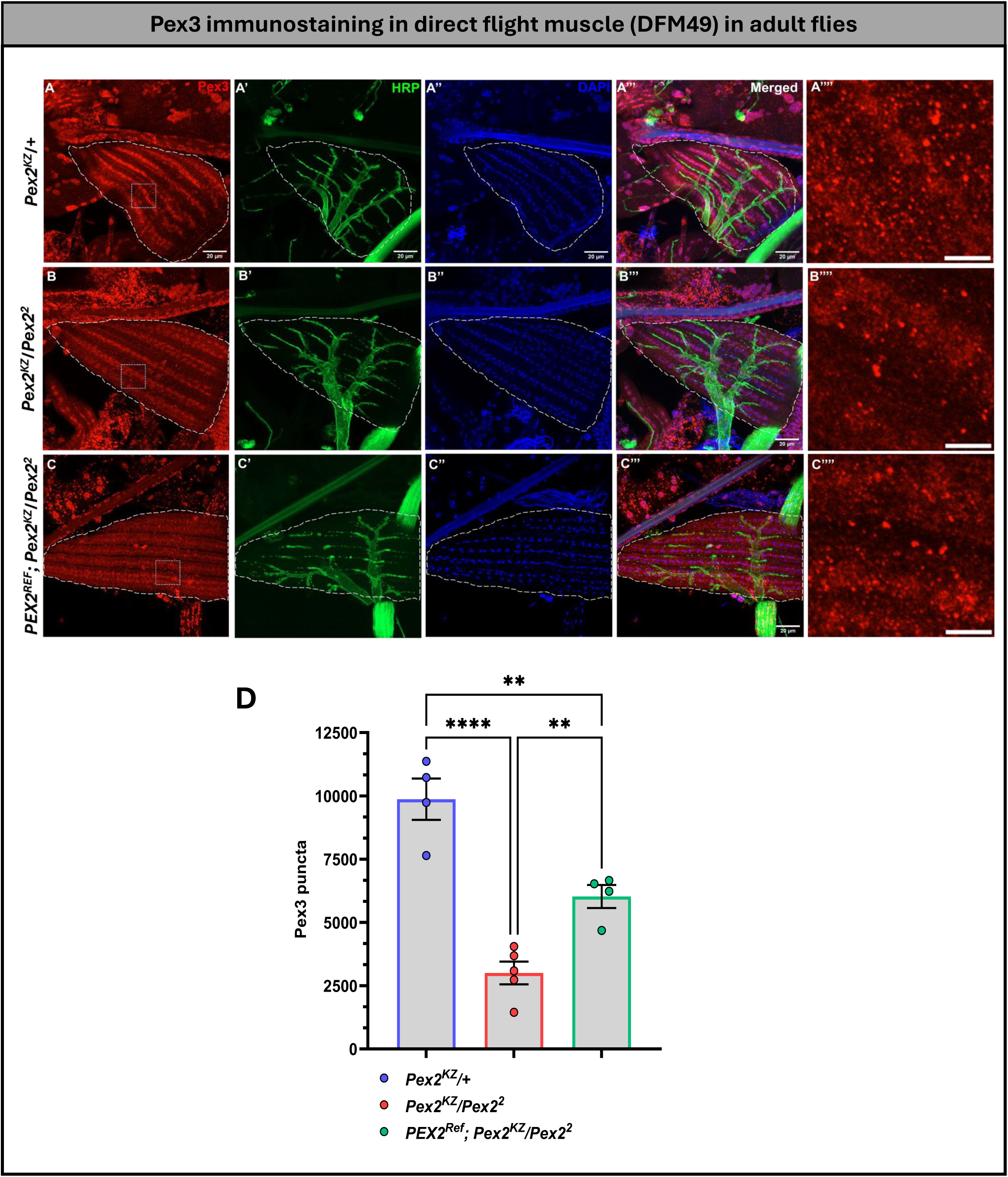
Human *PEX2* expression in *Pex2* null background significantly rescues peroxisome number in direct flight muscle (DFM49) in adult flies. (A, A’, A’’ & A’’’) represents control group; (B, B’, B’’ & B’’’) represents *Pex2* null flies & (C, C’, C’’ & C’’’) represents human rescue group. (A’’’’, B’’’ & C’’’’) illustrates the closure look at Pex3 puncta within the square box selected ROI for each respective group i.e., control, knockdown & rescue. Scale bar corresponds to 10 µm & 5 µm respectively. DFM49 marked with long dashed line. (D) Quantification of Pex3 puncta between the three genotypes. [**p<0.01; ****p<0.0001]

Similarly, we assessed the morphology of the indirect light muscle in *Pex16* flies. In our control flies, we observe Pex3 puncta intercalating the muscle fiber and HRP innervating the nerve fiber in 0-day-old flies (**Figure S5A**). In our *Pex16* mutant, we observed a dramatic reduction in Pex3 staining and thickening of the innervating nerve fiber (**Figure S5B**). However, the human *PEX16* transgene expression is able to rescue Pex3 staining and the nerve fiber thickening seen in the *Pex16* mutant (**Figure S5D**).

## Discussion

In this study, we generated “humanized” *Drosophila Pex2* and *Pex16* to the study the evolutionary conservation of the *PEX* genes across species and to assay the effect of human mutations^36,37^. In our study, we generated *KozakGAL4* alleles where we replaced the coding sequence of the fly gene with *GAL4*, generating a driver that allows expression of the human gene in its place^31^. Upon finding that these unique alleles effectively serve as nulls, we used the *KozakGAL4* lines to express human UAS-cDNAs of our targeted *PEX2* and *PEX16* gene and variants to assess rescue of the fly null phenotype. We began testing the impact of human proteins and their ability to rescue peroxisomal morphology defects. We found that *Pex2* and *Pex16* null mutants have a significantly reduced number of Pex3 puncta, but *PEX2^Ref^* and *PEX16^Ref^* can partially rescue the number of Pex3 puncta, demonstrating that the human protein can partially function in place of the fly protein *in vivo*. With these findings, *PEX2* and *PEX16* human reference constructs successfully restore peroxisomes, confirming the generation of humanized flies for *PEX2* and *PEX16*. It is remarkable to note that, for these two *Pex* genes, the human protein can participate in fly peroxisome biogenesis in distinct steps of the process, despite over 500 million years of evolution separating the fly and human genome.

In addition to humanization, another benefit of the *KozakGAL4* lines for *Pex2* and *Pex16 is* that we could see the endogenous expression pattern of these genes in the fly. While we would expect peroxisome biogenesis to be a general process, in this study we did not observe ubiquitous expression of either *Pex2* or *Pex16*. Moreover, there were differences in *Pex16* and *Pex2* expression patterns. We are relatively confident that our experiments mark the true expression of these genes because we find that human rescue is an independent confirmation for the respective *Pex* gene expression patterns. This is because the UAS human cDNA rescues the mutant phenotype while being expressed in a subset of cells. Since human rescue argues against the possibility that we do not see the full expression, we confirm that the expression patterns of *Pex2* and *Pex16* differ. This is somewhat surprising since peroxisome biogenesis requires both factors, as well as the other *Pex* genes.

Performing various behavior assays, we found that *PEX2^Ref^*and *PEX16^Ref^* were able to partially rescue *Pex2* and *Pex16* shortened lifespan, bang sensitivity, and defective climbing behavior. Interestingly, in addition to confirming human rescue, we also found that previously characterized “mild” or “atypical” PBD variants also produced a mild phenotype in the various assays. After utilizing rescue-based humanization of *Pex* genes, we found that our *PEX2^E55K^*and *PEX16^F332del^* variants display milder behavior phenotypes, compared to the other pathogenic variants. This finding corresponds to previous studies on mild PBD variants^21–24,38^, suggesting an allelic spectrum for *PEX2* and *PEX16*.

Based on previous studies, the four *PEX2* variants chosen range in clinical severity for classic PBD-ZSD^21,22,38–42^. The *PEX2^E55K^* variant has been identified in patients with mild PBD-ZSD in compound heterozygosity with the *PEX2^R1^*^19^*\** variant^21,22,38^. In our assays, we found that the *PEX2^E55K^* variant has a less severe phenotype compared to the other *PEX2* variants, having the best performance in lifespan analysis in females, the least severe bang-sensitive phenotype among the variants, and a slight decrease in climbing ability that is significantly better than the other variants. *PEX2^E55K^* never fully rescued the phenotypes, but except for male lifespan, it was consistently a hypomorph. Our other three *PEX2* variants have been characterized as pathogenic, but the clinical significance varies among patients. The homozygous *PEX2^W223*^* variant was identified in a patient with mild-PBD-ZSD^40,42^. While the *PEX2^W2^*^23^* flies did not perform as well in the various assays, it had a slightly better performance in viability assay and bang sensitivity than the *PEX2^R119*^* and *PEX2^C247R^*variants. The *PEX2^R119*^* variant has been identified in patients with severe and more mild cases of PBD-ZSD, where homozygous probands trend towards a more severe expression of the disease and patients with compound heterozygous genotype show milder or atypical presentations^22,39–41^. In flies, *PEX2^R119*^* had a more severe phenotype in behavior assays than the other two variants previously mentioned. Lastly, the presence of the homozygous *PEX2^C247R^* variant is also associated with a severe PBD-ZSD phenotype^40^. In flies, *PEX2^C247R^* also displayed the most severe phenotype in the behavior assays overall. With these findings, we propose an allelic severity spectrum with *PEX2^E55K^* being the least severe and *PEX2^C247R^* being the most severe (*PEX2^E55K^* < *PEX2^W223*^* < *PEX2^R119*^* < *PEX2^C247R^*) (**Table 1**).

**Table 1.** Clinical and Functional Severity of *PEX2* and *PEX16* gene mutations. *Pex2* and *Pex16* variant severity, considering clinical significance and severity, as well as outcomes observed from the study.

| *PEX2* Variant | Consequence | Clinical Significance | Clinical Severity | Outcomes Observed |
| --- | --- | --- | --- | --- |
| **E55K** | Missense | Likely Pathogenic | Mild PBD-ZSD in compound heterozygosity with R119* | - Viable F1 progeny<br>- Higher lifespan in females<br>- Less severe bang-sensitivity<br>- Slight climbing ability |
| **W223\*** | Nonsense | Pathogenic | Mild PBD-ZSD in homozygosity | - Somewhat viable F1 progeny<br>- Second less severe bang-sensitivity<br>- Decreased lifespan in males and females<br>- Severe climbing defect |
| **R119\*** | Nonsense | Pathogenic | - Severe PBD-ZSD in homozygosity<br>- Mild PBD-ZSD in compound heterozygosity with E55K<br>- Atypical PBD-ZSD in compound heterozygosity with 1-bp insertion in *PEX2* | - Semi-lethal F1 progeny<br>- Third less severe bang-sensitivity<br>- Decreased lifespan in males and females<br>- Severe climbing defect |
| **C247R** | Missense | Pathogenic | Severe PBD-ZSD in homozygosity | - Semi-lethal F1 progeny<br>- Most severe bang-sensitivity<br>- Decreased lifespan in males and females<br>- Severe climbing defect |

| *PEX16* Variant | Consequence | Clinical Significance | Clinical Severity | Outcomes Observed |
| --- | --- | --- | --- | --- |
| **F332del** | Inframe deletion | Likely Pathogenic | Atypical PBD-ZSD in homozygosity | - Viable F1 progeny<br>- Higher lifespan in females and males<br>- Less severe bang-sensitivity<br>- Climbing ability better than *PEX2* reference |
| **R176\*** | Nonsense | Pathogenic | Severe PBD-ZSD in homozygosity | - Semi-lethal F1 progeny<br>- Decreased lifespan in males and females<br>- Severe bang-sensitivity<br>- Severe climbing defect |

Additionally, we also found an allelic spectrum with our two *PEX16* variants. The *PEX16^F332del^*variant was reported by our group in a patient homozygous for the variant and with an atypical presentation of PBD-ZSD, having ataxia but normal peroxisomal biochemical testing in plasma^23,24^. In flies, we found that the *PEX16^F332del^* had completely viable F1 progeny, a lifespan similar to *PEX16^Ref^*, could rescue bang-sensitive, and even outperformed the human reference in the climbing assay at day 15. Alternatively, the homozygous *PEX16^R176*^*variant was seen in a patient who presented with a severe PBD-ZSD phenotype^43,44^. In flies, *PEX16^R176*^* was observed to have much more severe behavior phenotypes consistent with a null allele. *PEX16^R176*^* flies have semi-lethality in their F1 progeny, a decreased lifespan, severe bang sensitivity, and a severe climbing defect. With these findings, we propose an allelic severity spectrum with *PEX16^F332del^*being a mild variant and *PEX16^R176*^* being a severe variant (*PEX16^F332del^* < *PEX16^R176*^*) (**Table 1**).

Our data is informative for understanding the difference between mild PBD-ZSD with classic involvement of the retina and hearing versus the atypical ataxia phenotypes. The *PEX16^F332del^* variant has been associated with atypical ataxia with normal retinal and hearing findings. This atypical ataxia phenotype for *PEX16* has been characterized by other groups but lacks the classic features of PBD-ZSD, and its relationship to the classic spectrum has been unclear^46,47^. Our data shows that the ataxia allele behaves as a very weak hypomorphic allele in some contexts and is normal or even better than reference in others. This suggests that a certain level of very mild *PEX16* alleles can produce unique phenotypes that are not clinically recognizable as PBD-ZSD.

Severe alleles in *PEX* genes lead to severe clinical presentations, but in recent years, the mild presentations due to *PEX* mutations have expanded from phenotypes labeled as “infantile Refsum disease” to include atypical ataxia in *PEX16* and Heimler syndrome in *PEX1* and *PEX6*. How these mild conditions can differ so drastically is not yet understood. Using humanized flies to continue building an allelic spectrum as additional variants emerge has the great potential to fully distinguish the differences between these alleles. In the future, it will be interesting to determine how severe Heimler syndrome alleles alter the behavior and peroxisomal morphology in *Drosophila* compared to those studied here. *Drosophila* allows multiple *in vivo* assays for these variants to probe the pleiotropic effects of *PEX* genes on the neurological system.

## Methods

### Fly strains and maintenance

All flies were maintained at room temperature (21C), except when otherwise noted. Experiments were conducted at room temperature, as well.

### *Pex2* lines

*TI{KozakGAL4} Pex2 [CR70193-KO-kG4]* (labeled as *Pex2^KZ^*) (BDSC# 94997) were generated as described^31^. sgRNA target sites in *Pex2* UTRs are as follows: (TTTGTTTATATTCTTGCCTTTGG and GGTTCTGCGTGTCCTGAGTCGGG). Information about homology arms and the primers used to verify the insertions can be found at https://flypush.research.bcm.edu/pscreen/crimic/crimic.php.

The *Pex2^1^* and *Pex2^2^* lines were derived from imprecise excision of *(w[1118]; P{w[+mC] = E[g}pex2[HP35039]/TM3Sb[1]*. These were then backcrossed with 5 generations with

*yw:FRT80B* and studied as:

*yw ; FRT80B-Pex2^1^*(labeled as *Pex2^1^*)

*yw ; FRT80B-Pex2^2^*(labeled as *Pex2^2^*)

Additionally, a genomic deficiency line uncovering the *Pex2* locus was used as w118;Df(3L)BSC376/TM6C,Sb1 (labeled as *Pex2^Df^*)

For the generation of the human UAS-*PEX2* lines for Reference, E55K, R119X, W223X, C247R we obtained constructs which were codon-optimized for *Drosophila* and encoded the human protein from GeneART^TM^ (Thermo Fisher Scientific) (See **Supplemental Text**). These constructs were subcloned into the pUAST-attB vector using NotI and XhoI restriction sites. The UAS-constructs were then injected and inserted into the same genomic locus on the second chromosome (VK37 docking site) via φC31-mediated transgenesis^48^.

### *Pex16* lines

*TI{KozakGAL4} Pex16 [CR70194-KO-kG4]* (labeled as *Pex16^KZ^*) (BDSC# 94998) were generated as described^31^. sgRNA target sites in *Pex16* UTRs are as follows: ATAAAAATAATGAGGTGTTTCGG and TAGAGCGTTAGTATTCCCCTAGG).

Information about homology arms and the primers used to verify the insertions can be found at https://flypush.research.bcm.edu/pscreen/crimic/crimic.php.

The yw:Pex16^1^ line was obtained from Kenji Matsuno, derived from y^1^w^67c23^; P{w[+mC]y[+mDint2] = EPgy2}Pex16[EY05323].

*yw ; Pex16^1^*(labeled as *Pex16^1^*)

For the generation of the human UAS-*PEX16* lines for Reference, R176X and del955TCT (F332del), we obtained constructs which were codon-optimized for *Drosophila* and encoded the human protein from GeneART^TM^ (Thermo Fisher Scientific) (See **Supplemental Text**). These constructs were subcloned into the pUAST-attB vector using NotI and XhoI restriction sites. The UAS-constructs were then injected and inserted into the same genomic locus on the second chromosome (VK37 docking site) via φC31-mediated transgenesis^48^.

### Lifespan determination

Flies were collected under CO2 between 1 and 24 hours after eclosion. Male and female flies were separated and kept with 10 flies per vial at 25C. 100 flies of each line were collected, 50 females and 50 males. Fly food was changed every 3 days. The number of live flies was checked every 3 days until the last fly had died. A tally of the number of flies and their lifespan was kept, and the data was analyzed with a Kaplan-Meier survival curve.

### Bang sensitivity assay

Flies were kept without exposure to CO2 for at least 48 hours prior to the assay. Flies were vortexed for 10 seconds in an empty vial and the time it took to recover to normal behavior was recorded. Bang sensitivity was done for flies at 5 days, 10 days, and 15 days after eclosion.

### Climbing assay

Flies were kept without exposure to CO2 for at least 48 hours prior to the assay. Climbing assay was performed on flies at 5 days, 10 days, and 15 days after eclosion. Flies were dropped into an empty vial and timed to see how long it took to climb to the 8 cm mark within 60 seconds.

### Immunocytochemistry

Tissue samples were collected from wandering third instar larvae for body wall muscle and from the adult thorax for Direct flight muscle 49 (DFM49). Dissections were performed on Sylgard plates using 1x PBST (1% Tween 20 in 1x phosphate-buffered saline). The dissected larvae were fixed in 4% paraformaldehyde for 10 minutes at room temperature. After fixation, they were washed three times with PBST for 10 minutes each and then blocked with goat serum for 30 minutes.

The samples were incubated overnight at 4°C with primary antibodies, followed by three additional washes with PBST. Next, they were incubated overnight at 4°C with secondary antibodies and washed again three times with PBST. Finally, the samples were mounted on slides using Vectashield containing DAPI.

The primary antibody used was rabbit anti-Pex3 at a dilution of 1:500, (from the McNew lab at Rice University). The secondary antibody was donkey anti-rabbit IgG conjugated to Cy3 at a dilution of 1:1000. For immunostaining neuromuscular junctions in adult DFM49, we used Alexa Fluor 488 conjugated rabbit anti-horseradish peroxidase (HRP) at a dilution of 1:1000, which specifically recognizes the HRP epitope present on the surface of all *Drosophila* neurons.

### Fluorescence microscopy imaging

Imaging was performed using a Zeiss LSM800 Airyscan confocal microscope. Images were acquired with a Plan-Apochromat 40x/1.2 NA water immersion objective, using a frame size of 1024x1024 pixels. The excitation and emission wavelengths were as follows: ex405/em410-480 nm for DAPI, ex488/em493-570 nm for AF488, and ex561/em576-700 nm. The raw images were processed using the Airyscan Processing module in Zen 2.6 – Blue edition (Carl Zeiss Microscopy GmbH, Germany), applying the 2D SR processing option. The Airyscan filtering, utilizing a Wiener filter for deconvolution, was set to Standard.

### Image quantitative analysis

All fluorescence images were quantified and analyzed using the Surface module of Imaris v9.8.2 (Bitplane, Zurich, Switzerland). In this module, surfaces were generated for the fluorophore signal of interest, employing the background subtraction option and setting a lower area limit of 0.5 µm². This threshold was established to reduce background noise and minimize false positives during surface creation. Voxels outside the surface were masked and assigned a zero value. The number of surface puncta was then obtained from the statistical data generated for the created surface.

### Statistical analysis

Statistical analysis is completed using GraphPad Prism (Version 10.1.1). Continuous analysis is completed by Ordinary one-way ANOVA with multiple comparisons test, where differences between groups are quantified and a *p*-value less than 0.05 is considered significant.

## Acknowledgments

We acknowledge Carlos Bacino for introducing us to *PEX16* related ataxia. We also acknowledge the families affected by Peroxisomal disorders.

## Supplementary Figure Legend

**Figure S1** *Drosophila Pex16* mutants have shortened lifespan, are bang-sensitive, and have a climbing defect. (A) Schematic representation of fly *Pex16* gene along with one frameshift alleles (*Pex16^1^*) and *Pex16*-*KozakGAL4 (Pex16^KZ^*). (B) *Pex16* female lifespan assay shows that the *Pex16* mutants have a shorter lifespan compared to control lines (pink and purple). (C) *Pex16* male lifespan assay shows that the *Pex16* mutants have a shorter lifespan compared to control lines. D) *Pex16* null flies have a significant bang-sensitive phenotype (red) compared to controls (pink and purple) observed at 10 days after eclosion (DAE). (E) *Pex16* null flies have a significant climbing deficiency (red) compared to controls (pink and purple) observed at 10 days after eclosion. [* = *p*-value is less than 0.05. ** = *p*-value is less than 0.01. *** = *p*-value is less than 0.001. **** = *p*- value is less than 0.0001]

**Figure S2** Human UAS cDNA *PEX16* reference and variant lines. (A) Schematic representation of human *PEX16* gene. (B) Schematic representation of human PEX16 protein and variant locations. (C) *PEX16* variant table indicates the consequence of the change, pathogenicity prediction, clinical significance, clinical severity in homozygosity and heterozygosity, and conservation in *Drosophila.* (D) Indicates the observed/expected Mendelian ratio of the F1 generation of human *PEX16* variants, in a fly null background. (E) Assessment of the phenotype of indicated genotypes as lethal, viable, or semi-lethal.

**Figure S3** Human *PEX16* expression in *Pex16* null background larvae significantly rescues peroxisomes in 3^rd^ instar larva body wall 6 muscle. (A, A’ & A’’) represents control group; (B, B’, & B’’) represents *Pex16* null flies & (C, C’ & C’’) represents human rescue group. (D) Quantification of Pex3 positive puncta between the three genotypes. [**p=0.0011; ***p=0.001; ****p<0.0001]

**Figure S4** Rescue-based humanization of *Pex16*: Behavior Assays. (A) Lifespan analysis of *PEX16^Ref^ ^and^ ^Variant^* female flies, along with our *Pex16* null and control lines. (B) Lifespan analysis of *PEX16^Ref^ ^and^ ^Variant^* male flies, along with our *Pex16* null and control lines. (C) Bang sensitivity assay of *PEX16^Ref^ ^and^ ^Variant^* female flies, along with our *Pex16* null and control lines, at 10 days after eclosion (DAE). (D) Bang sensitivity assay of *PEX16^Ref^ ^and^ ^Variant^*female flies, along with our *Pex16* null and control lines, at 15 days after eclosion. (E-G) Climbing assay of *PEX16^Ref^ ^and^ ^Variant^*female flies, along with our *Pex16* null and control lines, at 5 days, 10 days, and 15 days after eclosion. [* = *p*-value is less than 0.05. ** = *p*-value is less than 0.01. *** = *p*-value is less than 0.001. **** = *p*- value is less than 0.0001]

**Figure S5** Human *PEX16* expression in *Pex16* null background significantly rescues peroxisome number in direct flight muscle (DFM49) in adult flies. (A, A’, A’’ & A’’’) represents control group; (B, B’, B’’ & B’’’) represents *Pex16* null flies & (C, C’, C’’ & C’’’) represents rescue group. (A’’’’, B’’’’ & C’’’’) illustrates the closure look at Pex3 puncta within the square box selected ROI for each respective group i.e., control, knockdown & rescue. Scale bar corresponds to 10 µm & 5 µm respectively. DFM49 marked with long dashed line.

