## Supplementary figures and images for "Distinguishing *PEX* gene variant severity for mild, severe, and atypical peroxisome biogenesis disorders in *Drosophila*"

### Figure S1

#### *Drosophila Pex16* mutants have shortened lifespan, are bang sensitive, and have a climbing defect

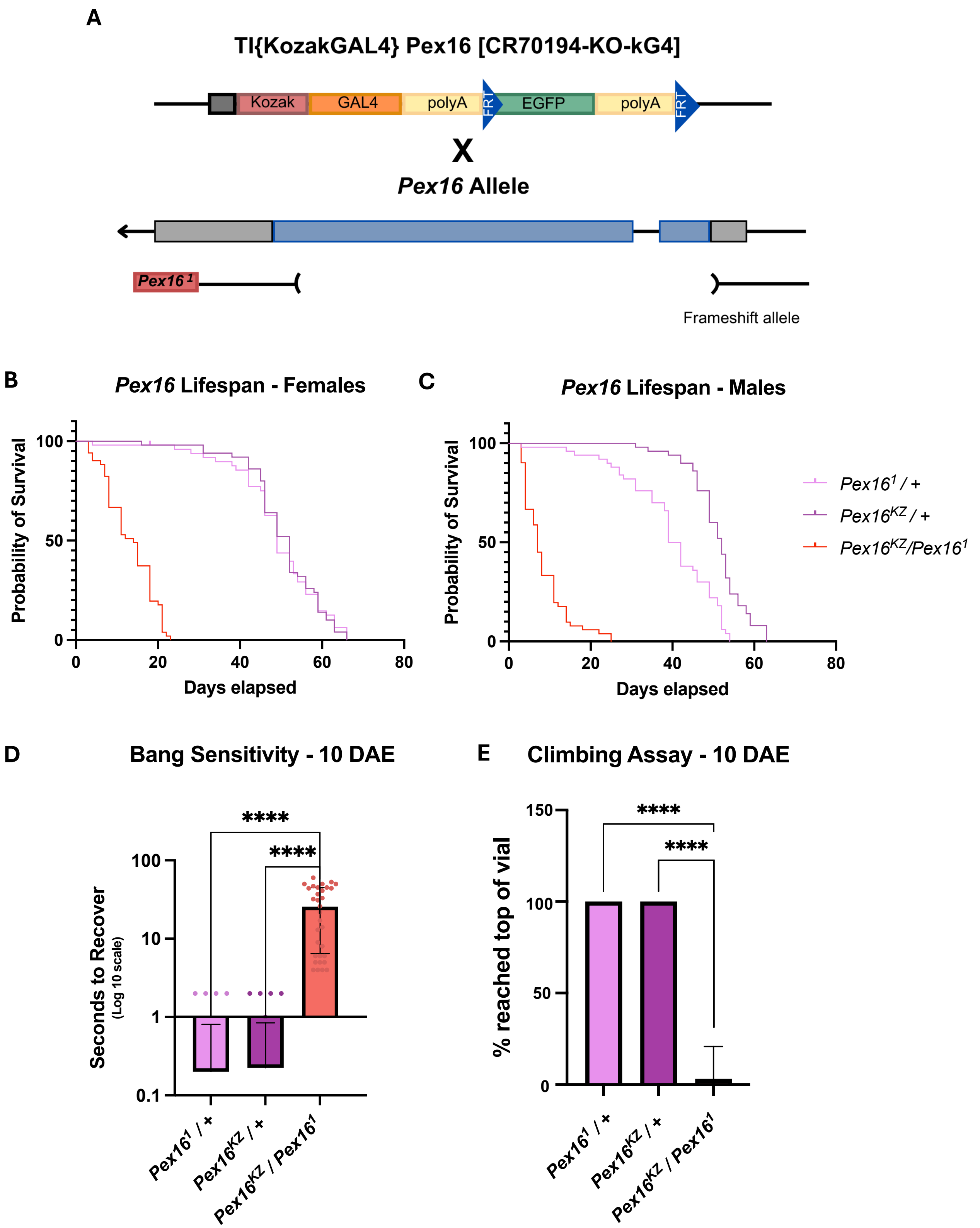

### Figure S3

Pex3 immunostaining in 3rd instar larva body wall 6 muscle

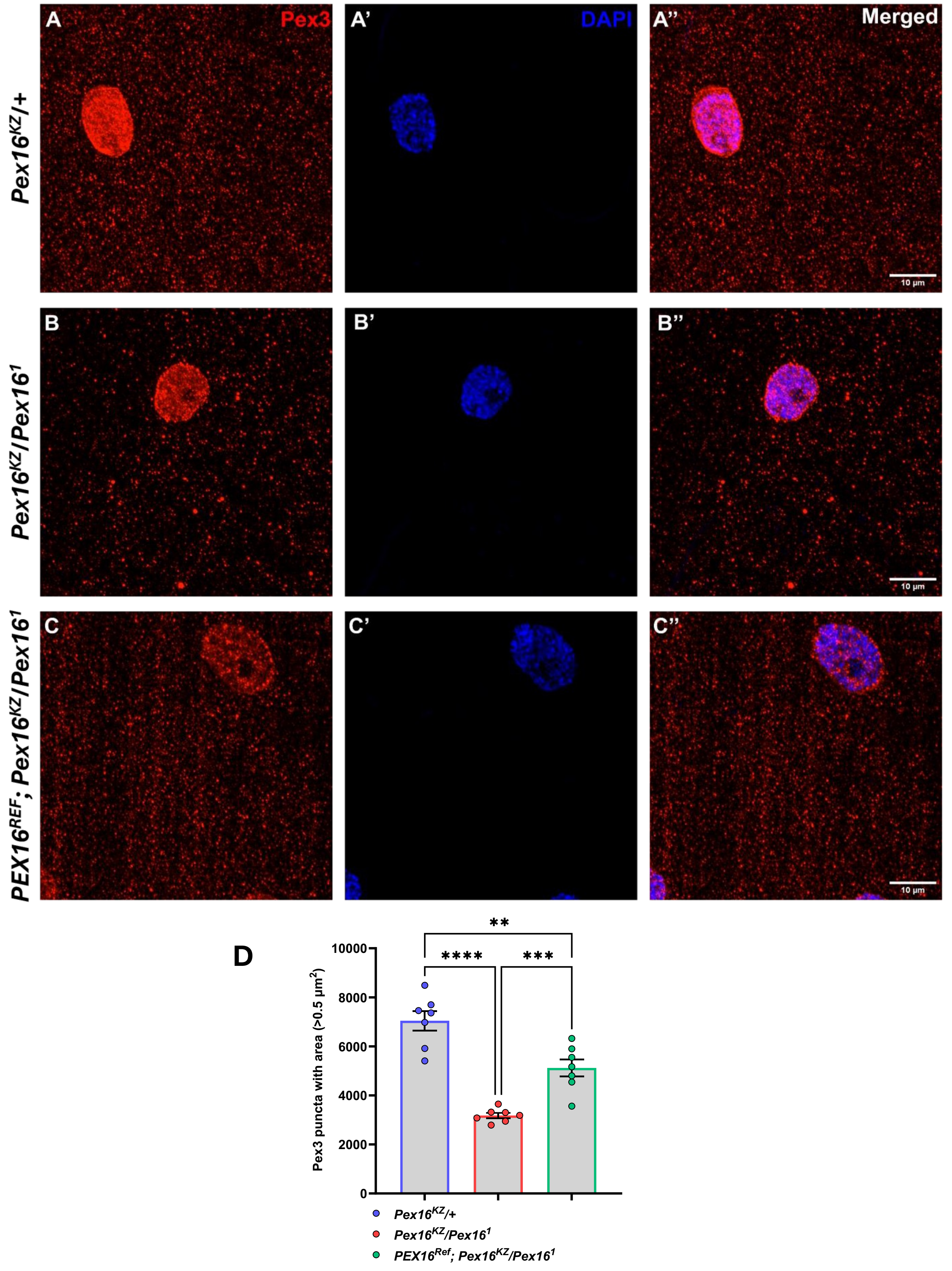

### Figure S4

#### Rescue-based humanization of *Pex16*: Behavior assays

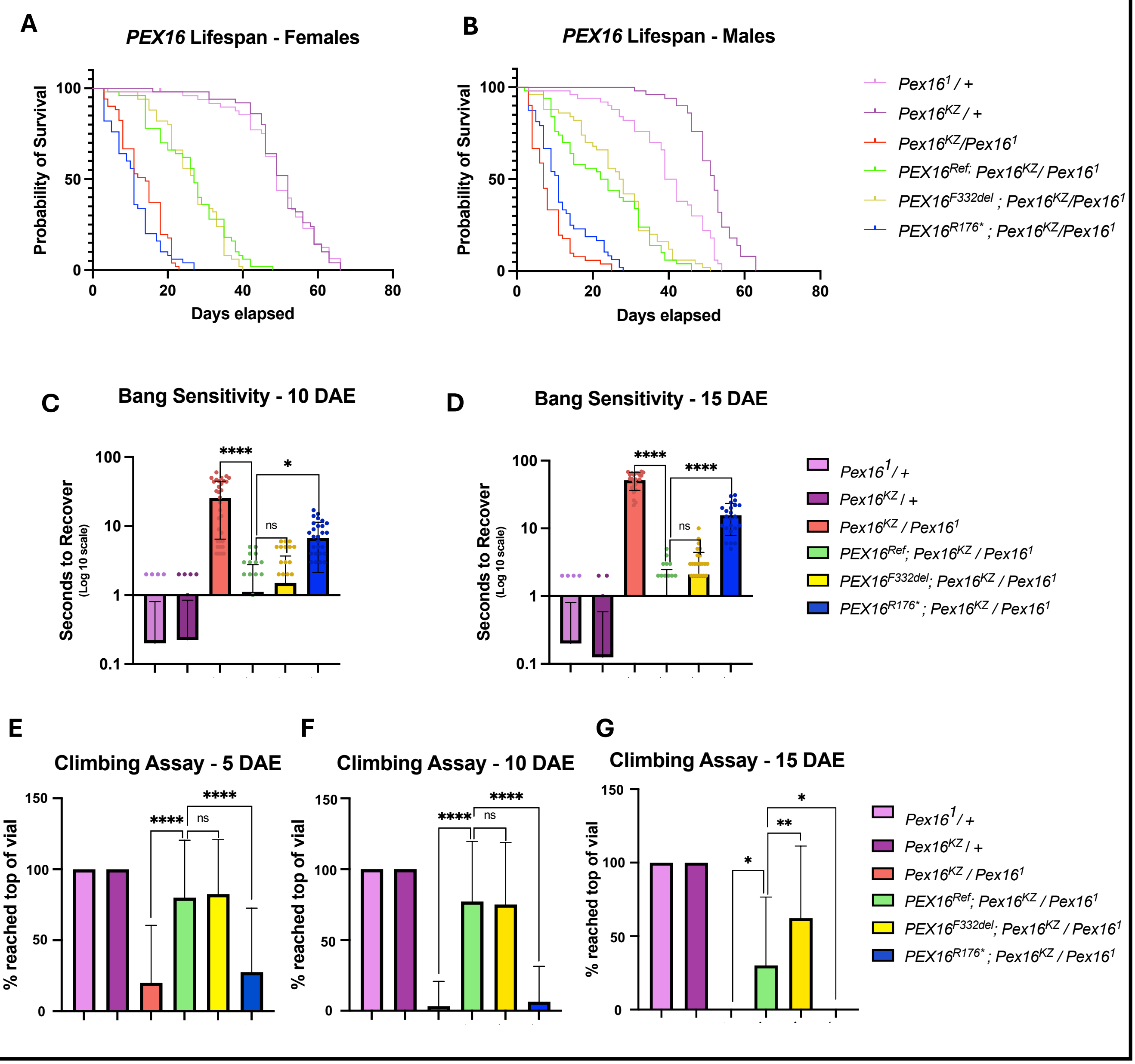

### Figure S5

#### Pex3 immunostaining in direct flight muscle (DFM49) in adult flies

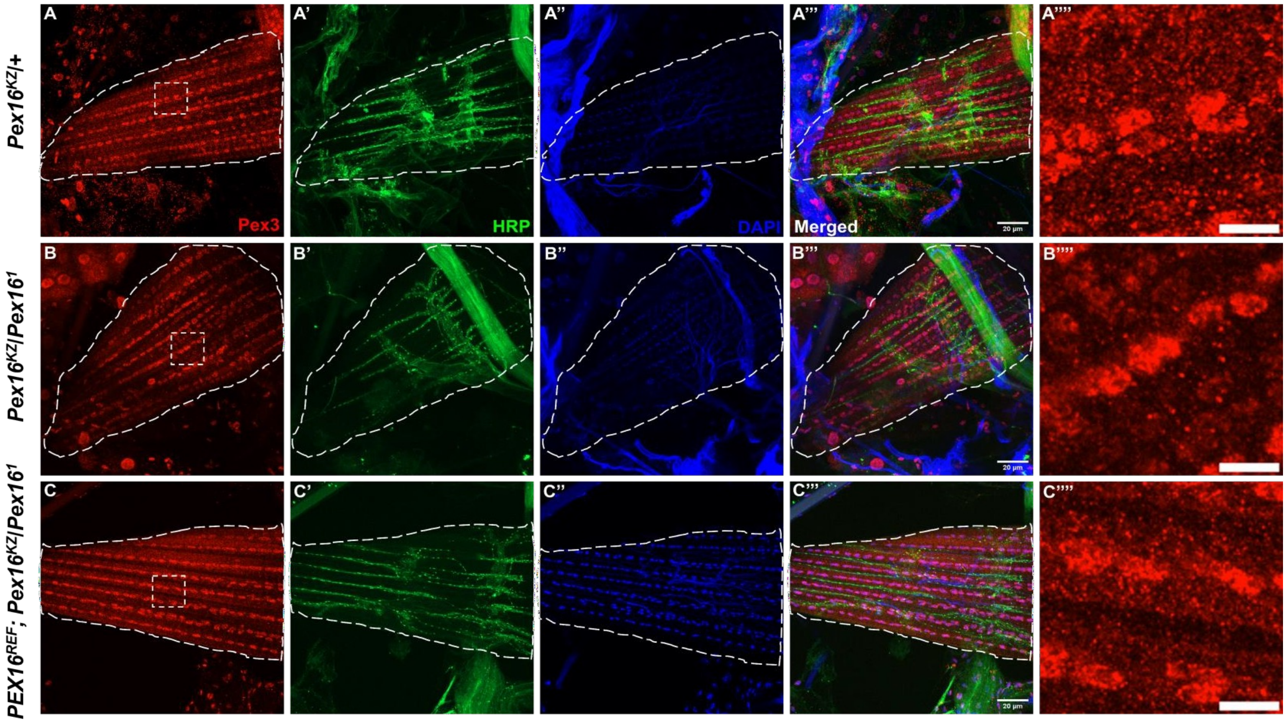
