## Supplementary material for "Distinguishing *PEX* gene variant severity for mild, severe, and atypical peroxisome biogenesis disorders in *Drosophila*": Figure S2

### Human UAS cDNA *PEX16* reference and variant lines

A

B

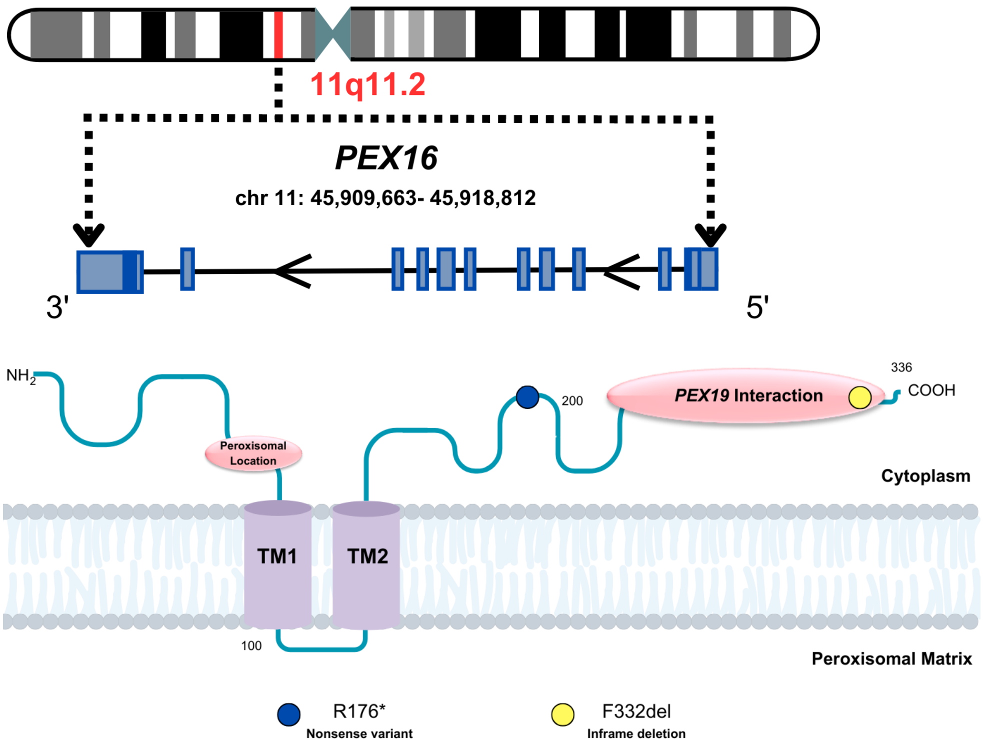


C

| *PEX16* Variant | Change | Consequence | CADD Score | Clinical Significance | Clinical Severity in Homozygosity | Clinical Severity in Heterozygosity | Conserved in *Drosophila*? |
|---|---|---|---|---|---|---|---|
| **R176*** | NM_004813.4 c.526C.T p.Arg176Ter | Nonsense | 37 | Pathogenic | **Severe PBD-ZSD** | Not seen | Yes |
| **F332del** | NM_0048813.4 c.995_997del p.Phe332del | Inframe deletion | - | Likely pathogenic | **Atypical PBD-ZSD**, presenting as ataxia | Not seen | Yes |

D

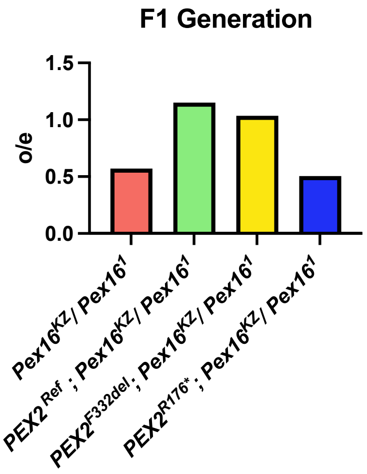


E

| Genotype | F1 Progeny |
|---|---|
| $Pex16^{1}$ / $Pex16^{KZ}$ | Semi-lethal |
| $PEX16^{Ref}$ ; $Pex16^{1}$ / $Pex16^{KZ}$ | Viable |
| $PEX16^{F332del}$ ; $Pex16^{1}$ / $Pex16^{KZ}$ | Viable |
| $PEX16^{R176*}$ ; $Pex16^{1}$ / $Pex16^{KZ}$ | Semi-lethal |
