## Supplemental text for "Distinguishing *PEX* gene variant severity for mild, severe, and atypical peroxisome biogenesis disorders in *Drosophila*"

**Sequence name / optimized for**

#### PEX2_Human_/ Drosophila melanogaster

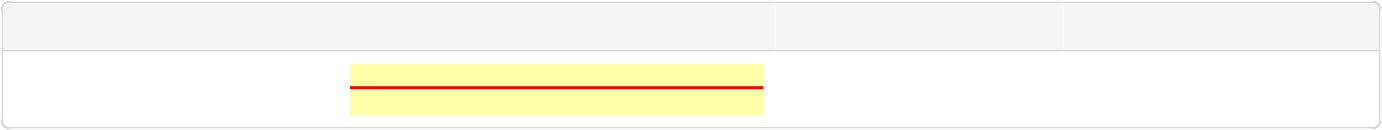

**ORF**

**Protected sites**

**Protected areas**

**Motifs to avoid**

13-1644 [ATG...TAG]

NotI [GCGGCCGC] XhoI [CTCGAG]

1-8 NotI [GCGGCCGC]

1645-1650 XhoI [CTCGAG]

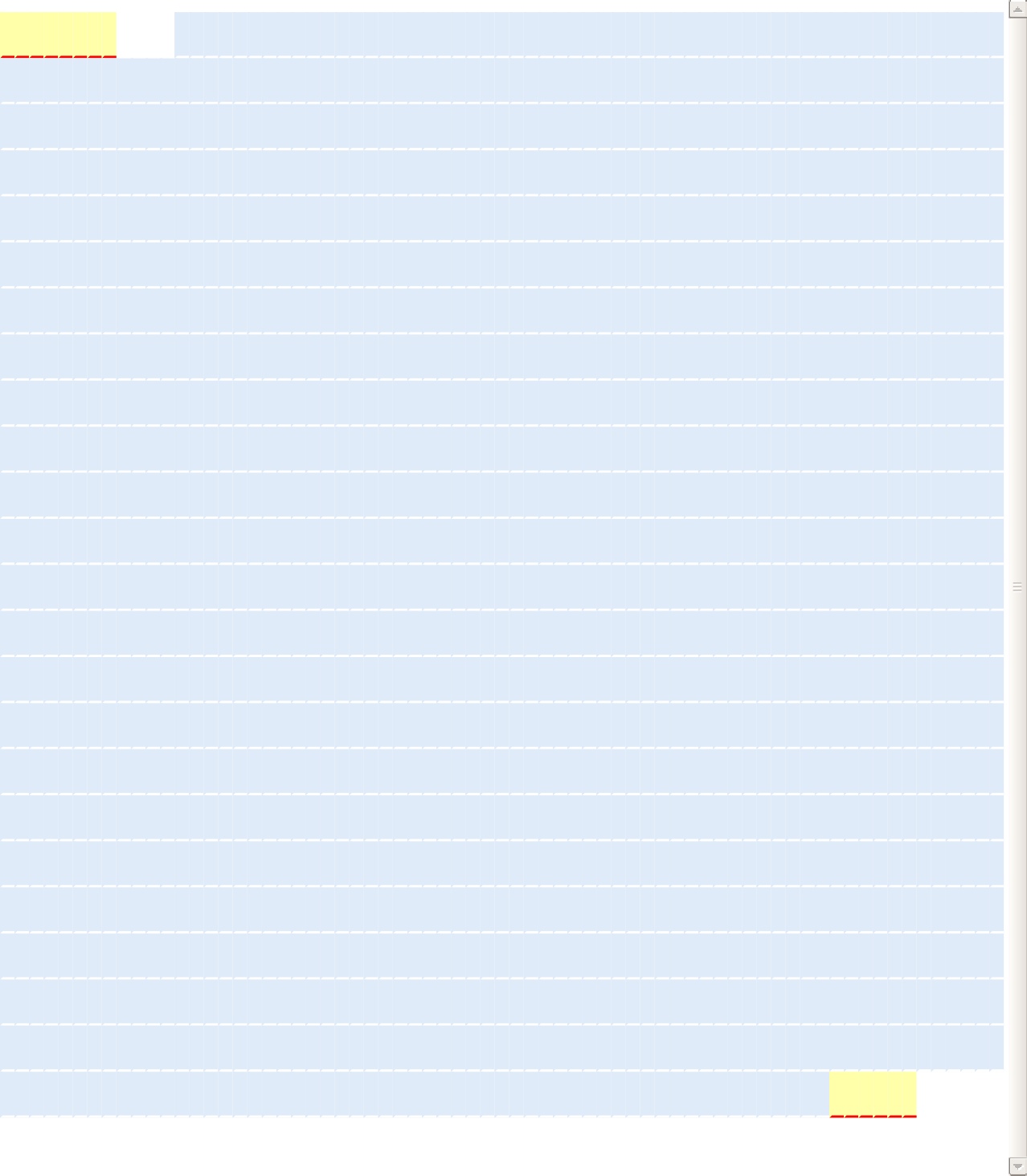
**M	A	S	R	K	E	N	A	K	S	A	N	R	V	L	R	I	S	Q**

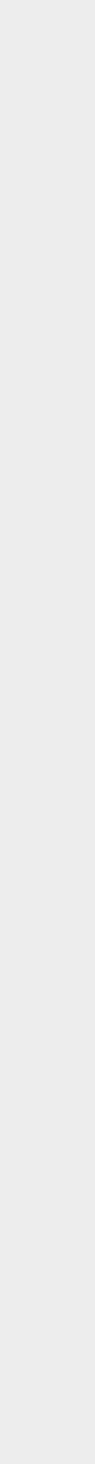

1.

70.

139.

208.

277.

346.

415.

484.

553.

622.

691.

760.

829.

898.

967.

1036.

1105.

1174.

1243.

1312.

1381.

1450.

1519.

1588.

G C G G C C G C C A A A A T G G C C A G C C G C A A G G A G A A T G C C A A G A G C G C C A A T C G C G T G C T G C G C A T C A G T C A G

**L	D	A**

**L	E	L	N	K	A	L**

**E	Q	L	V	W	S	Q	F	T	Q	C	F	H**

C T G G A T G C C C T G G A G C T G A A C A A G G C C C T G G A G C A G C T C G T G T G G T C C C A G T T C A C C C A G T G C T T C C A C

**G	F	K**

**P	G	L	L	A	R	F**

**E	P	E	V	K	A	C	L	W	V	F	L	W**

G G C T T C A A G C C A G G A C T G C T G G C C C G C T T T G A G C C C G A A G T G A A G G C C T G C C T G T G G G T G T T C C T G T G G

**R	F	T**

**I	Y	S	K	N	A	T**

**V	G	Q	S	V	L	N	I	K	Y	K	N	D**

C G C T T C A C C A T C T A C A G C A A G A A C G C C A C C G T G G G C C A G A G C G T G C T G A A C A T C A A G T A C A A G A A C G A C

**F	S	P**

**N	L	R	Y	Q	P	P**

**S	K	N	Q	K	I	W	Y	A	V	C	T**	**I**

T T C A G C C C C A A C C T G C G C T A C C A G C C C C C G A G C A A G A A C C A G A A G A T T T G G T A C G C C G T G T G C A C C A T C

**G	G	R**

**W	L	E	E	R	C	Y**

**D	L	F	R	N	H	H	L	A	S	F	G	K**

G G C G G A C G C T G G C T G G A G G A G C G C T G C T A C G A T C T G T T C C G C A A C C A C C A C C T G G C C A G C T T C G G C A A A

**V	K	Q**

**C	V	N	F	V	I	G**

**L	L	K	L	G	G	L	I	N	F	L	I	F**

G T G A A G C A G T G C G T G A A C T T C G T G A T C G G C C T G C T G A A G C T G G G C G G C C T G A T C A A C T T C C T G A T C T T C

**L	Q	R**

**G	K	F	A	T	L	T**

**E	R	L	L	G	I	H	S	V	F	C	K	P**

C T G C A G C G C G G C A A G T T C G C C A C C C T G A C C G A G C G C C T G C T G G G C A T T C A T A G C G T G T T C T G C A A G C C C

**Q	N**	**I**

**R	E	V	G	F	E	Y**

**M	N	R	E	L	L	W	H	G	F	A	E	F**

C A G A A C A T C C G C G A A G T G G G C T T C G A G T A C A T G A A C C G C G A G C T G C T G T G G C A C G G C T T C G C C G A G T T T

**L	I	F**

**L	L	P	L	I	N**	**V**

**Q	K	L	K	A	K	L	S	S	W	C	I	P**

C T G A T T T T C C T G C T G C C G C T G A T C A A C G T G C A G A A G C T G A A G G C C A A G C T G A G C A G C T G G T G C A T C C C A

**L	T	G**

**A	P	N	S	D	N	T**

**L	A	T	S	G	K	E	C	A	L	C	G	E**

C T G A C G G G A G C C C C C A A C A G C G A T A A C A C C C T G G C C A C C A G C G G A A A G G A G T G C G C C C T G T G C G G A G A G

**W	P	T**

**M	P	H	T	I	G	C**

**E	H	I	F	C	Y	F	C	A	K	S	S	F**

T G G C C A A C C A T G C C A C A C A C C A T T G G C T G C G A G C A C A T C T T C T G C T A C T T T T G C G C C A A G A G C A G C T T C

**L	F	D**

**V	Y	F	T	C	P	K**

**C	G	T	E	V	H	S	L	Q	P	L	K	S**

C T G T T C G A C G T G T A C T T C A C G T G C C C C A A G T G C G G C A C C G A G G T G C A C A G T C T G C A G C C A C T G A A G T C C

**G	I	E**

**M	S	E	V	N	A	L**

**V	S	K	G	E	E	L	F	T	G	V	V	P**

G G C A T C G A G A T G A G C G A A G T G A A C G C C C T G G T G T C C A A G G G C G A G G A G C T G T T T A C C G G C G T G G T G C C C

**I	L**	**V**

E **L	D	G	D	V	N**

**G	H	K	F	S	V	S	G	E	G	E	G	D**

A T T C T G G T G G A G C T G G A T G G C G A C G T G A A C G G C C A C A A G T T C A G C G T G T C C G G C G A G G G C G A G G G C G A C

**A	T	Y**

**G	K	L	T	L	K	F**

**I	C	T	T	G	K	L	P	V	P	W	P	T**

G C C A C C T A T G G A A A G C T G A C C C T G A A G T T C A T C T G C A C C A C C G G C A A G C T G C C C G T G C C A T G G C C A A C C

**L	V	T**

**T	L	T	Y	G	V	Q**

**C	F	S	R	Y	P	D	H	M	K	Q	H	D**

C T C G T G A C C A C G C T G A C C T A T G G C G T G C A G T G C T T C A G C C G C T A C C C C G A T C A C A T G A A G C A G C A C G A T

F **F	K**

**S	A	M	P	E	G	Y**

**V	Q	E	R	T	I	F	F	K	D	D	G	N**

T T C T T C A A G T C C G C C A T G C C C G A G G G C T A C G T G C A G G A G C G C A C C A T C T T T T T C A A G G A T G A C G G C A A C

**Y	K	T**

**R	A	E	V	K	F	E**

G **D	T	L	V	N	R	I	E	L	K	G**	**I**

T A C A A G A C C C G C G C C G A A G T G A A G T T C G A G G G C G A T A C C C T C G T G A A C C G C A T C G A G C T G A A G G G C A T C

**D	F	K**

**E	D	G	N	I	L	G**

H **K	L	E	Y	N	Y	N	S	H	N	V	Y**

G A T T T C A A G G A G G A T G G A A A C A T C C T G G G C C A C A A G C T G G A G T A C A A C T A C A A C A G C C A C A A C G T G T A C

I **M	A**

**D	K	Q	K	N	G**	**I**

**K	V	N	F	K	I	R	H	N	I	E	D	G**

A T C A T G G C C G A C A A G C A G A A G A A C G G C A T C A A A G T G A A C T T C A A G A T T C G C C A C A A C A T C G A G G A T G G C

**S	V	Q**

**L	A	D	H	Y	Q	Q**

**N	T	P	I	G	D	G	P	V	L	L	P	D**

A G C G T G C A G C T G G C C G A C C A C T A C C A G C A G A A C A C C C C C A T C G G A G A T G G C C C C G T G C T G C T G C C C G A T

**N	H	Y**

**L	S	T	Q	S	A	L**

**S	K	D	P	N	E	K	R	D	H	M	V	L**

A A C C A C T A C C T G A G T A C C C A G A G C G C C C T G A G C A A G G A T C C C A A C G A G A A G C G C G A C C A C A T G G T G C T G

**L	E	F**

**V	T	A	A	G	I	T**

L **G	M	D	E	L	Y	K	***

C T G G A G T T T G T G A C C G C C G C C G G C A T T A C C C T G G G C A T G G A T G A G C T G T A C A A G T A G C T C G A G

The histograms show the percentage of sequence codons which fall into a certain quality class. The quality value of the most frequently used codon for a given amino acid in the desired expression system is set to 100, the remaining codons are scaled accordingly (see also Sharp, P.M., Li, W.H., Nucleic Acids Res. 15 (3),1987).

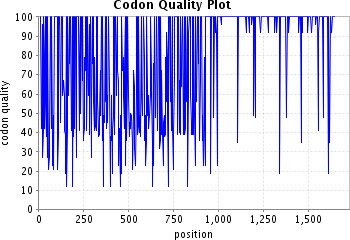

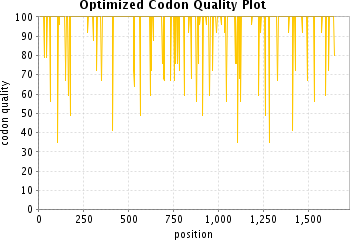

The plots show the quality of the used codon at the indicated codon position.

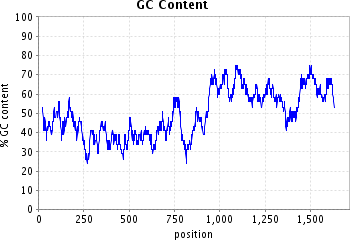

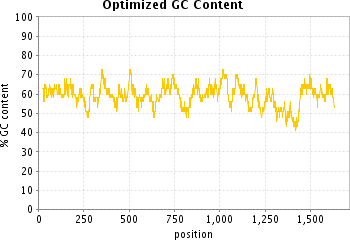

The plots show the GC content in a 40 bp window centered at the indicated nucleotide position.

**Sequence name / optimized for**

#### PEX2_E55K/ Drosophila melanogaster

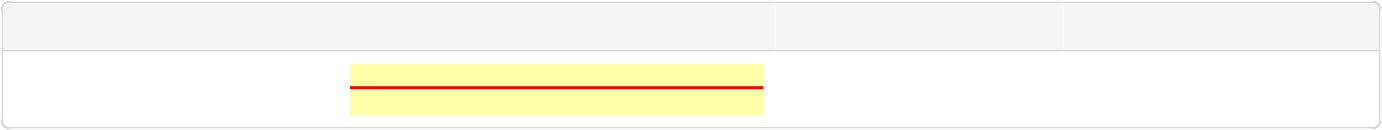

**ORF**

**Protected sites**

**Protected areas**

**Motifs to avoid**

13-1644 [ATG...TAG]

NotI [GCGGCCGC] XhoI [CTCGAG]

1-8 NotI [GCGGCCGC]

1645-1650 XhoI [CTCGAG]

M
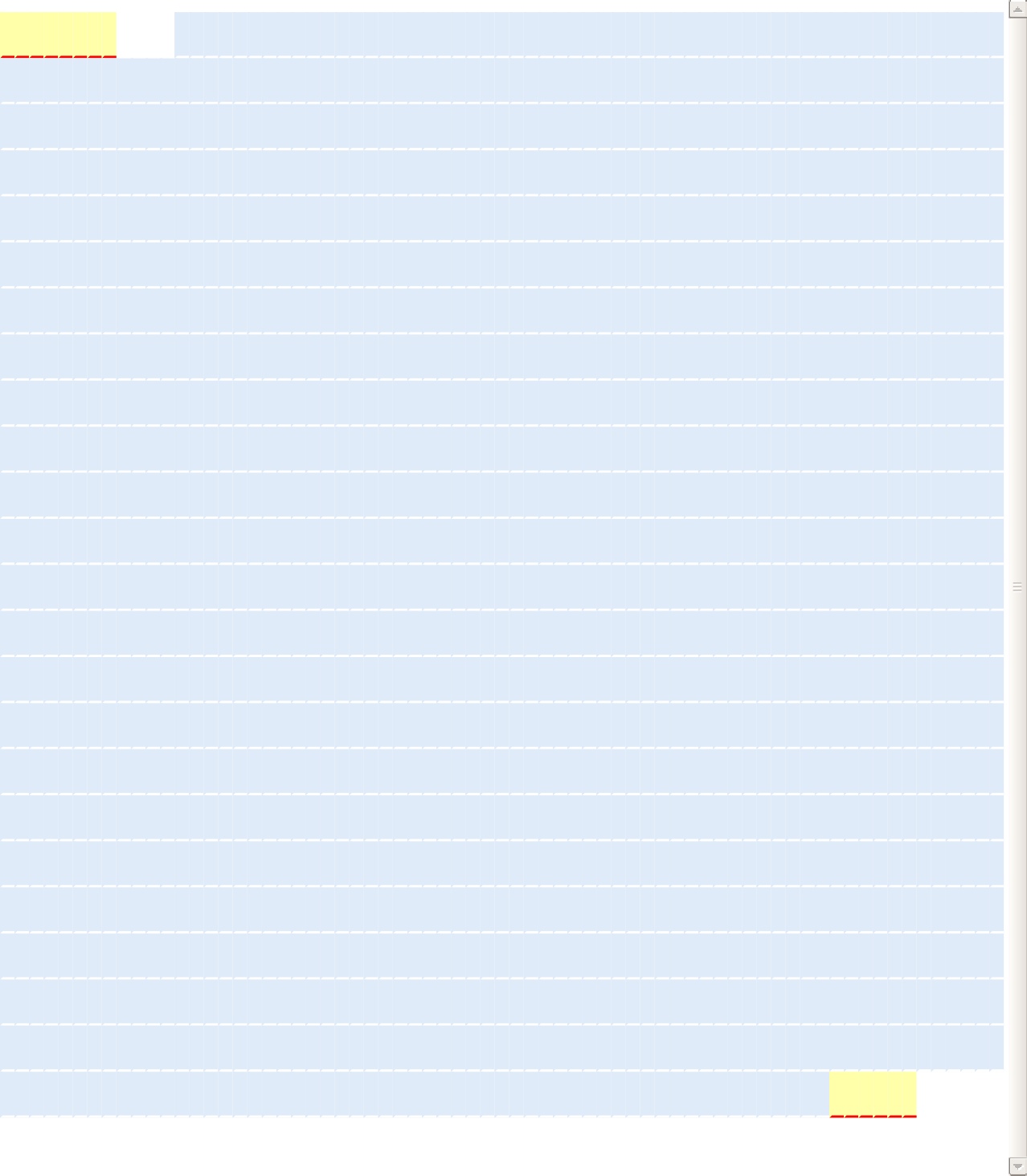
**A	S	R	K	E	N	A	K	S	A	N	R	V	L	R	I	S	Q**

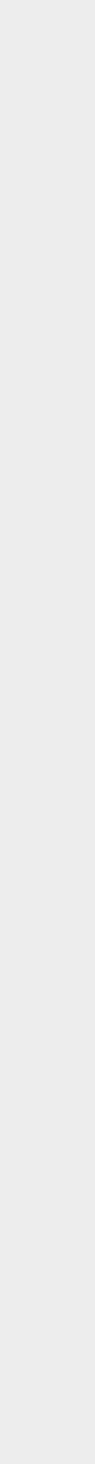

1.

70.

139.

208.

277.

346.

415.

484.

553.

622.

691.

760.

829.

898.

967.

1036.

1105.

1174.

1243.

1312.

1381.

1450.

1519.

1588.

G C G G C C G C C A A A A T G G C C A G C C G C A A G G A G A A T G C C A A G A G C G C C A A T C G C G T G C T G C G C A T C A G T C A G

**L	D	A**

**L	E	L	N	K	A	L**

**E	Q	L	V	W	S	Q	F	T	Q	C	F	H**

C T G G A T G C C C T G G A G C T G A A C A A G G C C C T G G A G C A G C T C G T G T G G T C C C A G T T C A C C C A G T G C T T C C A C

**G	F	K**

**P	G	L	L	A	R	F**

**E	P	K	V	K	A	C	L	W	V	F	L	W**

G G C T T C A A G C C A G G A C T G C T G G C C C G C T T C G A G C C C A A A G T G A A G G C C T G C C T G T G G G T G T T C C T G T G G

**R	F	T**

**I	Y	S	K	N	A	T**

**V	G	Q	S	V	L	N	I	K	Y	K	N	D**

C G C T T C A C C A T C T A C A G C A A G A A C G C C A C C G T G G G C C A G A G C G T G C T G A A C A T C A A G T A C A A G A A C G A C

**F	S	P**

**N	L	R	Y	Q	P	P**

**S	K	N	Q	K	I	W	Y	A	V	C	T**	**I**

T T C A G C C C C A A C C T G C G C T A C C A G C C C C C G A G C A A G A A C C A G A A G A T T T G G T A C G C C G T G T G C A C C A T C

**G	G	R**

**W	L	E	E	R	C	Y**

**D	L	F	R	N	H	H	L	A	S	F	G	K**

G G C G G A C G C T G G C T G G A G G A G C G C T G C T A C G A T C T G T T C C G C A A C C A C C A C C T G G C C A G C T T C G G C A A A

**V	K	Q**

**C	V	N	F	V	I	G**

**L	L	K	L	G	G	L	I	N	F	L	I	F**

G T G A A G C A G T G C G T G A A C T T C G T G A T C G G C C T G C T G A A G C T G G G C G G C C T G A T C A A C T T C C T G A T C T T C

**L	Q	R**

**G	K	F	A	T	L	T**

**E	R	L	L	G	I	H	S	V	F	C	K	P**

C T G C A G C G C G G C A A G T T C G C C A C C C T G A C C G A G C G C C T G C T G G G C A T T C A T A G C G T G T T C T G C A A G C C C

**Q	N**	**I**

**R	E	V	G	F	E	Y**

**M	N	R	E	L	L	W	H	G	F	A	E	F**

C A G A A C A T C C G C G A A G T G G G C T T C G A G T A C A T G A A C C G C G A G C T G C T G T G G C A C G G C T T C G C C G A G T T T

**L	I	F**

**L	L	P	L	I	N**	**V**

**Q	K	L	K	A	K	L	S	S	W	C	I	P**

C T G A T T T T C C T G C T G C C G C T G A T C A A C G T G C A G A A G C T G A A G G C C A A G C T G A G C A G C T G G T G C A T C C C A

**L	T	G**

**A	P	N	S	D	N	T**

**L	A	T	S	G	K	E	C	A	L	C	G	E**

C T G A C G G G A G C C C C C A A C A G C G A T A A C A C C C T G G C C A C C A G C G G A A A G G A G T G C G C C C T G T G C G G A G A G

**W	P	T**

**M	P	H	T	I	G	C**

**E	H	I	F	C	Y	F	C	A	K	S	S	F**

T G G C C A A C C A T G C C A C A C A C C A T T G G C T G C G A G C A C A T C T T C T G C T A C T T T T G C G C C A A G A G C A G C T T C

**L	F	D**

**V	Y	F	T	C	P	K**

C **G	T	E	V	H	S	L	Q	P	L	K	S**

C T G T T C G A C G T G T A C T T C A C G T G C C C C A A G T G C G G C A C C G A G G T G C A C A G T C T G C A G C C A C T G A A G T C C

**G	I	E**

**M	S	E	V	N	A	L**

**V	S	K	G	E	E	L	F	T	G	V	V	P**

G G C A T C G A G A T G A G C G A A G T G A A C G C C C T G G T G T C C A A G G G C G A G G A G C T G T T T A C C G G C G T G G T G C C C

**I	L**	**V**

E **L	D	G	D	V	N**

**G	H	K	F	S	V	S	G	E	G	E	G	D**

A T T C T G G T G G A G C T G G A T G G C G A C G T G A A C G G C C A C A A G T T C A G C G T G T C C G G C G A G G G C G A G G G C G A C

**A	T	Y**

**G	K	L	T	L	K	F**

**I	C	T	T	G	K	L	P	V	P	W	P	T**

G C C A C C T A T G G A A A G C T G A C C C T G A A G T T C A T C T G C A C C A C C G G C A A G C T G C C C G T G C C A T G G C C A A C C

**L	V	T**

**T	L	T	Y	G	V	Q**

**C	F	S	R	Y	P	D	H	M	K	Q	H	D**

C T C G T G A C C A C G C T G A C C T A T G G C G T G C A G T G C T T C A G C C G C T A C C C C G A T C A C A T G A A G C A G C A C G A T

F **F	K**

**S	A	M	P	E	G	Y**

**V	Q	E	R	T	I	F	F	K	D	D	G	N**

T T C T T C A A G T C C G C C A T G C C C G A G G G C T A C G T G C A G G A G C G C A C C A T C T T T T T C A A G G A T G A C G G C A A C

**Y	K	T**

**R	A	E	V	K	F	E**

G **D	T	L	V	N	R	I	E	L	K	G**	**I**

T A C A A G A C C C G C G C C G A A G T G A A G T T C G A G G G C G A T A C C C T C G T G A A C C G C A T C G A G C T G A A G G G C A T C

**D	F	K**

**E	D	G	N	I	L	G**

H **K	L	E	Y	N	Y	N	S	H	N	V	Y**

G A T T T C A A G G A G G A T G G A A A C A T C C T G G G C C A C A A G C T G G A G T A C A A C T A C A A C A G C C A C A A C G T G T A C

I **M	A**
D **K	Q	K	N	G**	**I**

**K	V	N	F	K	I	R	H	N	I	E	D	G**

A T C A T G G C C G A C A A G C A G A A G A A C G G C A T C A A A G T G A A C T T C A A G A T T C G C C A C A A C A T C G A G G A T G G C

**S	V	Q**

**L	A	D	H	Y	Q	Q**

**N	T	P	I	G	D	G	P	V	L	L	P	D**

A G C G T G C A G C T G G C C G A C C A C T A C C A G C A G A A C A C C C C C A T C G G A G A T G G C C C C G T G C T G C T G C C C G A T

**N	H	Y**

**L	S	T	Q	S	A	L**

**S	K	D	P	N	E	K	R	D	H	M	V	L**

A A C C A C T A C C T G A G T A C C C A G A G C G C C C T G A G C A A G G A T C C C A A C G A G A A G C G C G A C C A C A T G G T G C T G

**L	E	F**

**V	T	A	A	G	I	T**

L **G	M	D	E	L	Y	K	***

C T G G A G T T T G T G A C C G C C G C C G G C A T T A C C C T G G G C A T G G A T G A G C T G T A C A A G T A G C T C G A G

The histograms show the percentage of sequence codons which fall into a certain quality class. The quality value of the most frequently used codon for a given amino acid in the desired expression system is set to 100, the remaining codons are scaled accordingly (see also Sharp, P.M., Li, W.H., Nucleic Acids Res. 15 (3),1987).

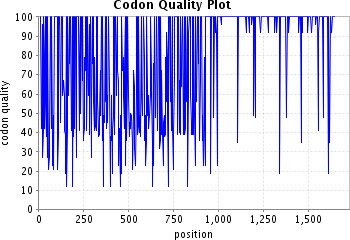

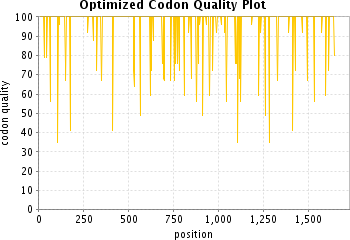

The plots show the quality of the used codon at the indicated codon position.

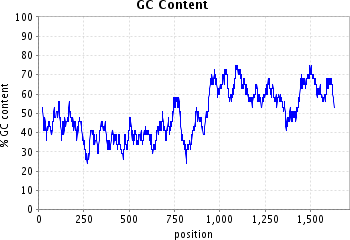

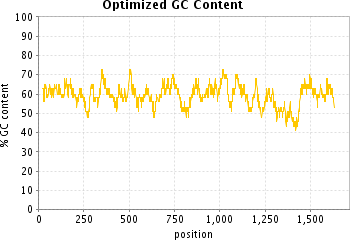

The plots show the GC content in a 40 bp window centered at the indicated nucleotide position.

**Sequence name / optimized for**

#### PEX2_R119X/ Drosophila melanogaster

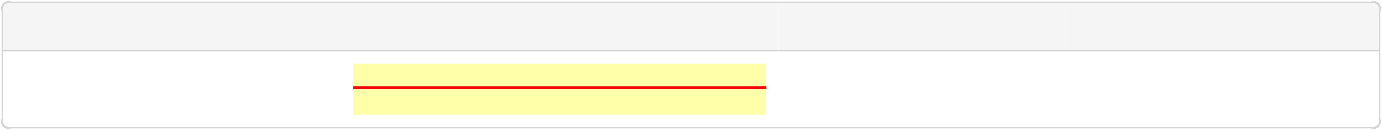

**ORF**

**Protected sites**

**Protected areas**

**Motifs to avoid**

13-1083 [ATG...TAG]

NotI [GCGGCCGC] XhoI [CTCGAG]

1-8 NotI [GCGGCCGC]

1084-1089 XhoI [CTCGAG]

M
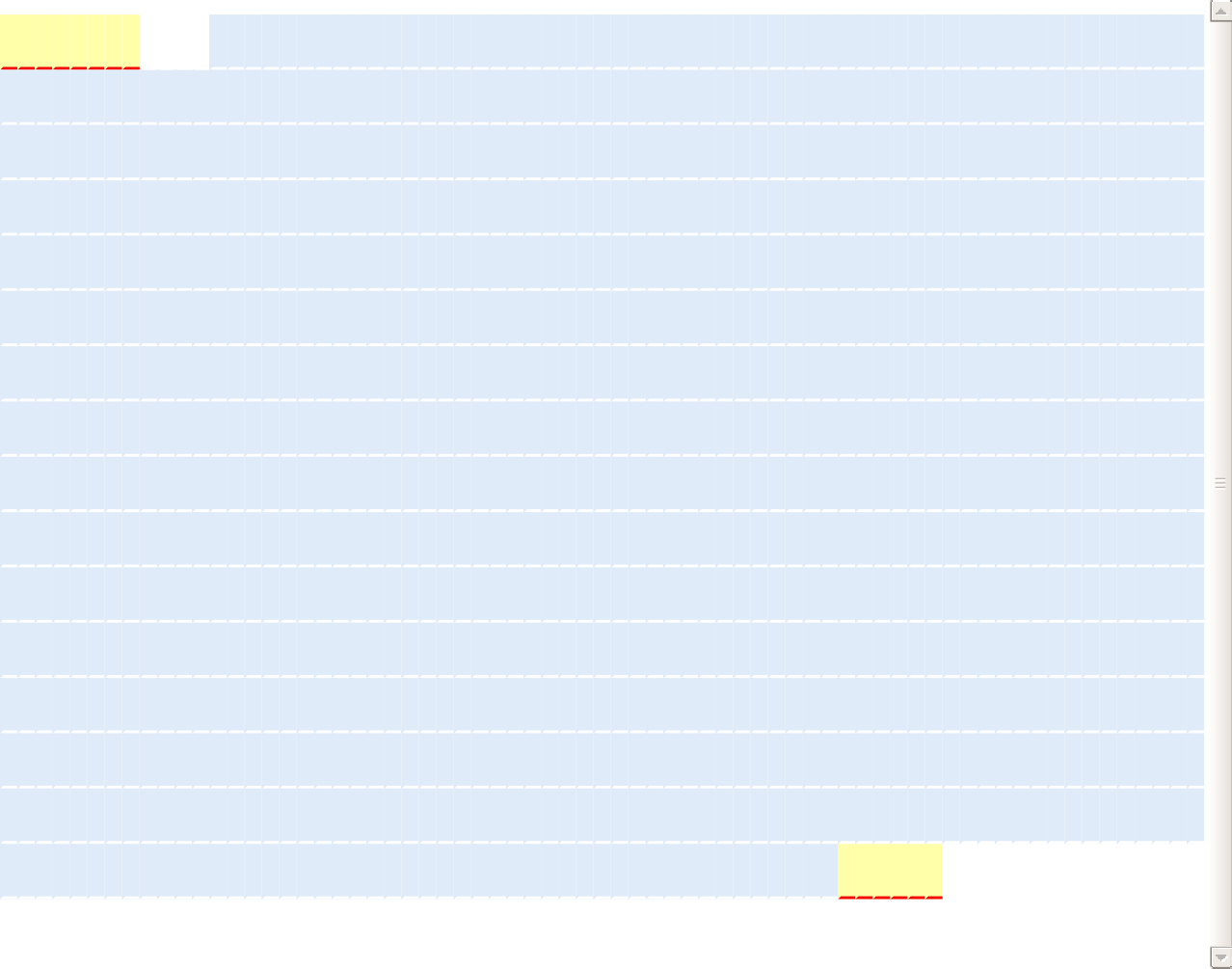
**A	S	R	K	E	N	A	K	S	A	N	R	V	L	R	I	S	Q**

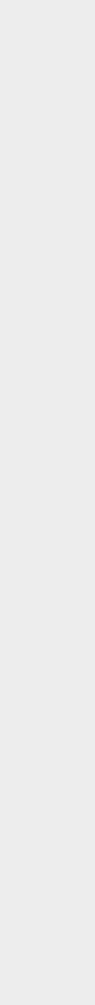

1.

70.

139.

208.

277.

346.

415.

484.

553.

622.

691.

760.

829.

898.

967.

1036.

G C G G C C G C C A A A A T G G C C A G C C G C A A G G A G A A T G C C A A G A G C G C C A A T C G C G T G C T G C G C A T C A G T C A G

**L	D	A**

**L	E	L	N	K	A	L**

E **Q	L	V	W	S	Q	F	T	Q	C	F	H**

C T G G A T G C C C T G G A G C T G A A C A A G G C C C T G G A G C A G C T C G T G T G G T C C C A G T T C A C C C A G T G C T T C C A C

**G	F	K**

**P	G	L	L	A	R	F**

**E	P	E	V	K	A	C	L	W	V	F	L	W**

G G C T T C A A G C C A G G A C T G C T G G C C C G C T T T G A G C C C G A A G T G A A G G C C T G C C T G T G G G T G T T C C T G T G G

**R	F	T**

**I	Y	S	K	N	A	T**

**V	G	Q	S	V	L	N	I	K	Y	K	N	D**

C G C T T C A C C A T C T A C A G C A A G A A C G C C A C C G T G G G C C A G A G C G T G C T G A A C A T C A A G T A C A A G A A C G A C

**F	S	P**

**N	L	R	Y	Q	P	P**

**S	K	N	Q	K	I	W	Y	A	V	C	T**	**I**

T T C A G C C C C A A C C T G C G C T A C C A G C C C C C G A G C A A G A A C C A G A A G A T T T G G T A C G C C G T G T G C A C C A T C

**G	G	R**

**W	L	E	E	V	S	K**

**G	E	E	L	F	T	G	V	V	P	I	L**	**V**

G G C G G A C G C T G G C T G G A G G A G G T G T C C A A G G G C G A G G A G C T G T T T A C C G G C G T G G T G C C C A T T C T G G T G

**E	L	D**

G **D	V	N	G	H	K**
F **S	V	S	G	E	G	E	G	D	A	T	Y**

G A G C T G G A T G G C G A C G T G A A C G G C C A C A A G T T C A G C G T G T C C G G C G A G G G C G A G G G C G A C G C C A C C T A T

G **K	L**

**T	L	K	F	I	C	T**

**T	G	K	L	P	V	P	W	P	T	L	V	T**

G G A A A G C T G A C C C T G A A G T T C A T C T G C A C C A C C G G C A A G C T G C C C G T G C C A T G G C C A A C C C T C G T G A C C

**T	L	T**

**Y	G	V	Q	C	F	S**

**R	Y	P	D	H	M	K	Q	H	D	F	F	K**

A C C C T G A C C T A T G G C G T G C A G T G C T T C A G C C G C T A C C C C G A T C A C A T G A A G C A G C A C G A T T T C T T C A A G

**S	A	M**

P **E	G	Y	V	Q	E**

**R	T	I	F	F	K	D	D	G	N	Y	K	T**

T C C G C C A T G C C C G A G G G C T A C G T G C A G G A G C G C A C C A T C T T T T T C A A G G A T G A C G G C A A C T A C A A G A C C

**R	A	E**

**V	K	F	E	G	D	T**

**L	V	N	R	I	E	L	K	G	I	D	F	K**

C G C G C C G A A G T G A A G T T C G A G G G C G A T A C C C T C G T G A A C C G C A T C G A G C T G A A G G G C A T C G A T T T C A A G

**E	D	G**

**N	I	L	G	H	K	L**

**E	Y	N	Y	N	S	H	N	V	Y	I	M	A**

G A G G A T G G A A A C A T C C T G G G C C A C A A G C T G G A G T A C A A C T A C A A C A G C C A C A A C G T G T A C A T C A T G G C C

**D	K	Q**

**K	N	G	I	K	V	N**

**F	K	I	R	H	N	I	E	D	G	S	V	Q**

G A C A A G C A G A A G A A C G G C A T C A A A G T G A A C T T C A A G A T T C G C C A C A A C A T C G A G G A T G G C A G C G T G C A G

**L	A	D**

H **Y	Q	Q	N	T	P**
I **G	D	G	P	V	L	L	P	D	N	H	Y**

C T G G C C G A T C A C T A C C A G C A G A A C A C C C C A A T C G G C G A C G G C C C A G T G C T G C T G C C C G A T A A C C A T T A C

L **S	T**

Q **S	A	L	S	K	D**

**P	N	E	K	R	D	H	M	V	L	L	E	F**

C T G A G C A C C C A G A G C G C C C T G A G C A A G G A T C C C A A C G A G A A G C G C G A C C A C A T G G T G C T G C T G G A G T T T

**V	T	A**

**A	G	I	T	L	G	M**

**D	E	L	Y	K	***

G T G A C C G C C G C C G G C A T T A C C C T G G G C A T G G A T G A G C T G T A C A A G T A G C T C G A G

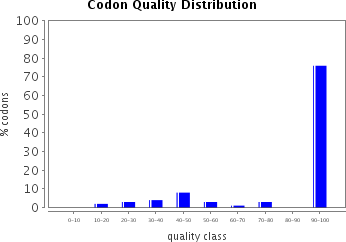

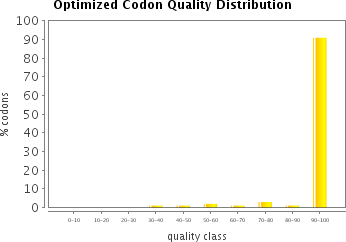

The histograms show the percentage of sequence codons which fall into a certain quality class. The quality value of the most frequently used codon for a given amino acid in the desired expression system is set to 100, the remaining codons are scaled accordingly (see also Sharp, P.M., Li, W.H., Nucleic Acids Res. 15 (3),1987).

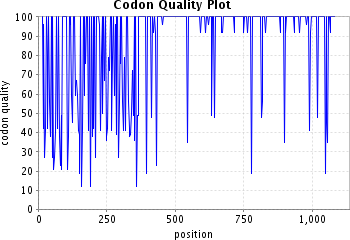

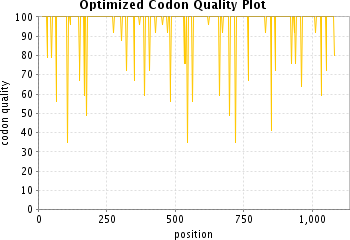

The plots show the quality of the used codon at the indicated codon position.

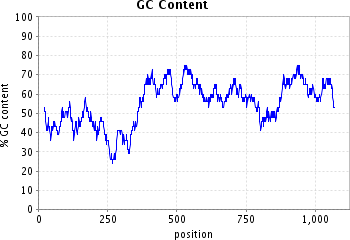

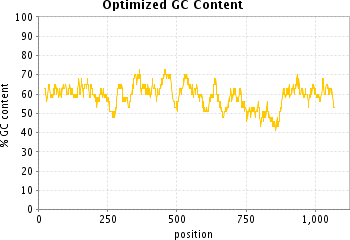

The plots show the GC content in a 40 bp window centered at the indicated nucleotide position.

**Sequence name / optimized for**

#### PEX2_W223X/ Drosophila melanogaster

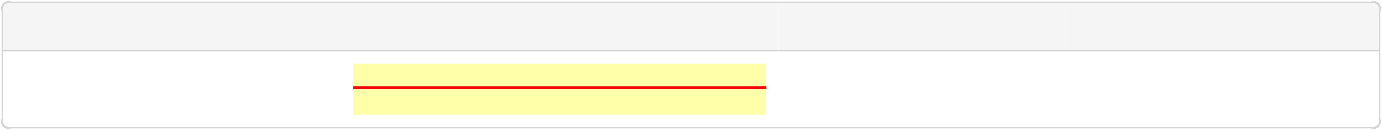

**ORF**

**Protected sites**

**Protected areas**

**Motifs to avoid**

13-1395 [ATG...TAG]

NotI [GCGGCCGC] XhoI [CTCGAG]

1-8 NotI [GCGGCCGC]

1396-1401 XhoI [CTCGAG]

M
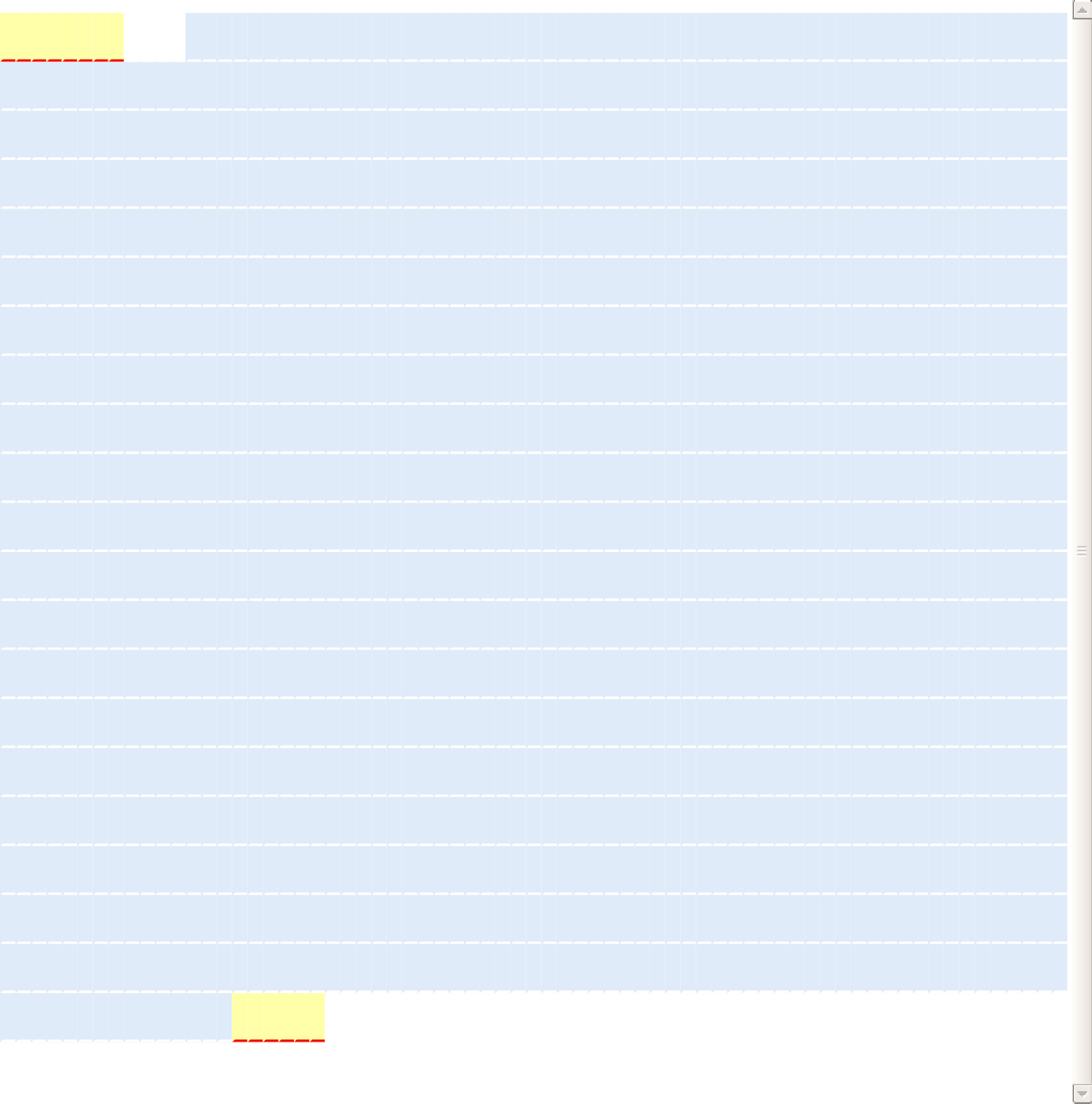
**A	S	R	K	E	N	A	K	S	A	N	R	V	L	R	I	S	Q**

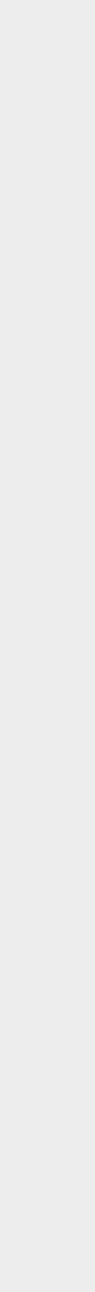

1.

70.

139.

208.

277.

346.

415.

484.

553.

622.

691.

760.

829.

898.

967.

1036.

1105.

1174.

1243.

1312.

1381.

G C G G C C G C C A A A A T G G C C A G C C G C A A G G A G A A T G C C A A G A G C G C C A A T C G C G T G C T G C G C A T C A G T C A G

**L	D	A**

**L	E	L	N	K	A	L**

**E	Q	L	V	W	S	Q	F	T	Q	C	F	H**

C T G G A T G C C C T G G A G C T G A A C A A G G C C C T G G A G C A G C T C G T G T G G T C C C A G T T C A C C C A G T G C T T C C A C

**G	F	K**

**P	G	L	L	A	R	F**

**E	P	E	V	K	A	C	L	W	V	F	L	W**

G G C T T C A A G C C A G G A C T G C T G G C C C G C T T T G A G C C C G A A G T G A A G G C C T G C C T G T G G G T G T T C C T G T G G

R **F	T**

**I	Y	S	K	N	A	T**

**V	G	Q	S	V	L	N	I	K	Y	K	N	D**

C G C T T C A C C A T C T A C A G C A A G A A C G C C A C C G T G G G C C A G A G C G T G C T G A A C A T C A A G T A C A A G A A C G A C

**F	S	P**

**N	L	R	Y	Q	P	P**

**S	K	N	Q	K	I	W	Y	A	V	C	T**	**I**

T T C A G C C C C A A C C T G C G C T A C C A G C C C C C G A G C A A G A A C C A G A A G A T T T G G T A C G C C G T G T G C A C C A T C

**G	G	R**

**W	L	E	E	R	C	Y**

D **L	F	R	N	H	H	L	A	S	F	G	K**

G G C G G A C G C T G G C T G G A G G A G C G C T G C T A C G A T C T G T T C C G C A A C C A C C A C C T G G C C A G C T T C G G C A A A

**V	K	Q**

**C	V	N	F	V	I	G**

**L	L	K	L	G	G	L	I	N	F	L	I	F**

G T G A A G C A G T G C G T G A A C T T C G T G A T C G G C C T G C T G A A G C T G G G C G G C C T G A T C A A C T T C C T G A T C T T C

**L	Q	R**

**G	K	F	A	T	L	T**

E **R	L	L	G	I	H	S	V	F	C	K	P**

C T G C A G C G C G G C A A G T T C G C C A C C C T G A C C G A G C G C C T G C T G G G C A T T C A T A G C G T G T T C T G C A A G C C C

**Q	N**	**I**

**R	E	V	G	F	E	Y**

**M	N	R	E	L	L	W	H	G	F	A	E	F**

C A G A A C A T C C G C G A A G T G G G C T T C G A G T A C A T G A A C C G C G A G C T G C T G T G G C A C G G C T T C G C C G A G T T T

**L	I	F**

**L	L	P	L	I	N**	**V**

**Q	K	L	K	A	K	L	S	S	V	S	K	G**

C T G A T T T T C C T G C T G C C G C T G A T C A A C G T G C A G A A G C T G A A G G C C A A G C T G A G C A G C G T G T C C A A G G G C

**E	E	L**

F **T	G	V	V	P**	**I**

**L	V	E	L	D	G	D	V	N	G	H	K	F**

G A G G A G C T G T T T A C C G G C G T G G T G C C C A T T C T G G T G G A G C T G G A T G G C G A C G T G A A C G G C C A C A A G T T C

S **V	S**
G **E	G	E	G	D	A**

**T	Y	G	K	L	T	L	K	F	I	C	T	T**

T C C G T G T C C G G C G A G G G C G A G G G C G A C G C C A C C T A T G G A A A G C T G A C C C T G A A G T T C A T C T G C A C C A C C

**G	K	L**

**P	V	P	W	P	T	L**

**V	T	T	L	T	Y	G	V	Q	C	F	S	R**

G G C A A G C T G C C C G T G C C A T G G C C A A C C C T C G T G A C C A C G C T G A C C T A T G G C G T G C A G T G C T T C A G C C G C

**Y	P	D**

H **M	K	Q	H	D	F**

**F	K	S	A	M	P	E	G	Y	V	Q	E	R**

T A C C C C G A T C A C A T G A A G C A G C A C G A T T T C T T C A A G T C C G C C A T G C C C G A G G G C T A C G T G C A G G A G C G C

T **I	F**

**F	K	D	D	G	N	Y**

**K	T	R	A	E	V	K	F	E	G	D	T	L**

A C C A T C T T T T T C A A G G A T G A C G G C A A C T A C A A G A C C C G C G C C G A A G T G A A G T T C G A G G G C G A T A C C C T C

**V	N	R**

**I	E	L	K	G	I	D**

**F	K	E	D	G	N	I	L	G	H	K	L	E**

G T G A A C C G C A T C G A G C T G A A G G G C A T C G A T T T C A A G G A G G A T G G A A A C A T C C T G G G C C A C A A G C T G G A G

**Y	N	Y**

**N	S	H	N	V	Y**	**I**

**M	A	D	K	Q	K	N	G	I	K	V	N	F**

T A C A A C T A C A A C A G C C A C A A C G T G T A C A T C A T G G C C G A C A A G C A G A A G A A C G G C A T C A A A G T G A A C T T C

**K	I	R**

**H	N	I	E	D	G	S**

**V	Q	L	A	D	H	Y	Q	Q	N	T	P**	**I**

A A G A T T C G C C A C A A C A T C G A G G A T G G C A G C G T G C A G C T G G C C G A C C A C T A C C A G C A G A A C A C C C C C A T C

**G	D	G**

**P	V	L	L	P	D	N**

**H	Y	L	S	T	Q	S	A	L	S	K	D	P**

G G A G A T G G C C C C G T G C T G C T G C C C G A T A A C C A C T A C C T G A G T A C C C A G A G C G C C C T G A G C A A G G A T C C C

**N	E	K**

**R	D	H	M	V	L	L**

**E	F	V	T	A	A	G	I	T	L	G	M	D**

A A C G A G A A G C G C G A C C A C A T G G T G C T G C T G G A G T T T G T G A C C G C C G C C G G C A T T A C C C T G G G C A T G G A T

**E	L	Y	K	***

G A G C T G T A C A A G T A G C T C G A G

The histograms show the percentage of sequence codons which fall into a certain quality class. The quality value of the most frequently used codon for a given amino acid in the desired expression system is set to 100, the remaining codons are scaled accordingly (see also Sharp, P.M., Li, W.H., Nucleic Acids Res. 15 (3),1987).

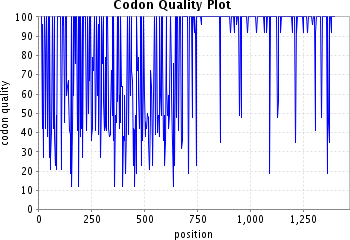

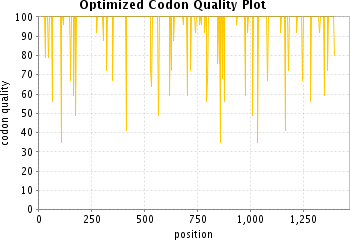

The plots show the quality of the used codon at the indicated codon position.

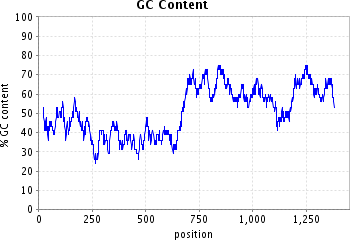

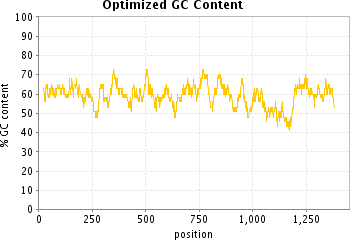

The plots show the GC content in a 40 bp window centered at the indicated nucleotide position.

**Sequence name / optimized for**

#### PEX2_C247R/ Drosophila melanogaster

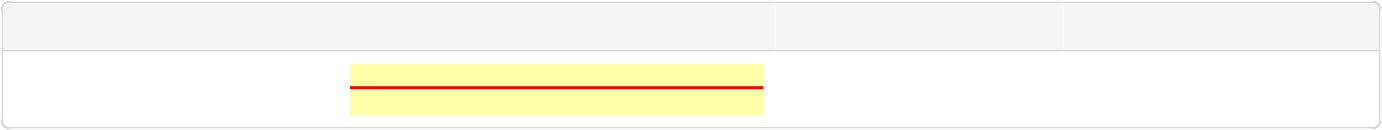

**ORF**

**Protected sites**

**Protected areas**

**Motifs to avoid**

13-1644 [ATG...TAG]

NotI [GCGGCCGC] XhoI [CTCGAG]

1-8 NotI [GCGGCCGC]

1645-1650 XhoI [CTCGAG]

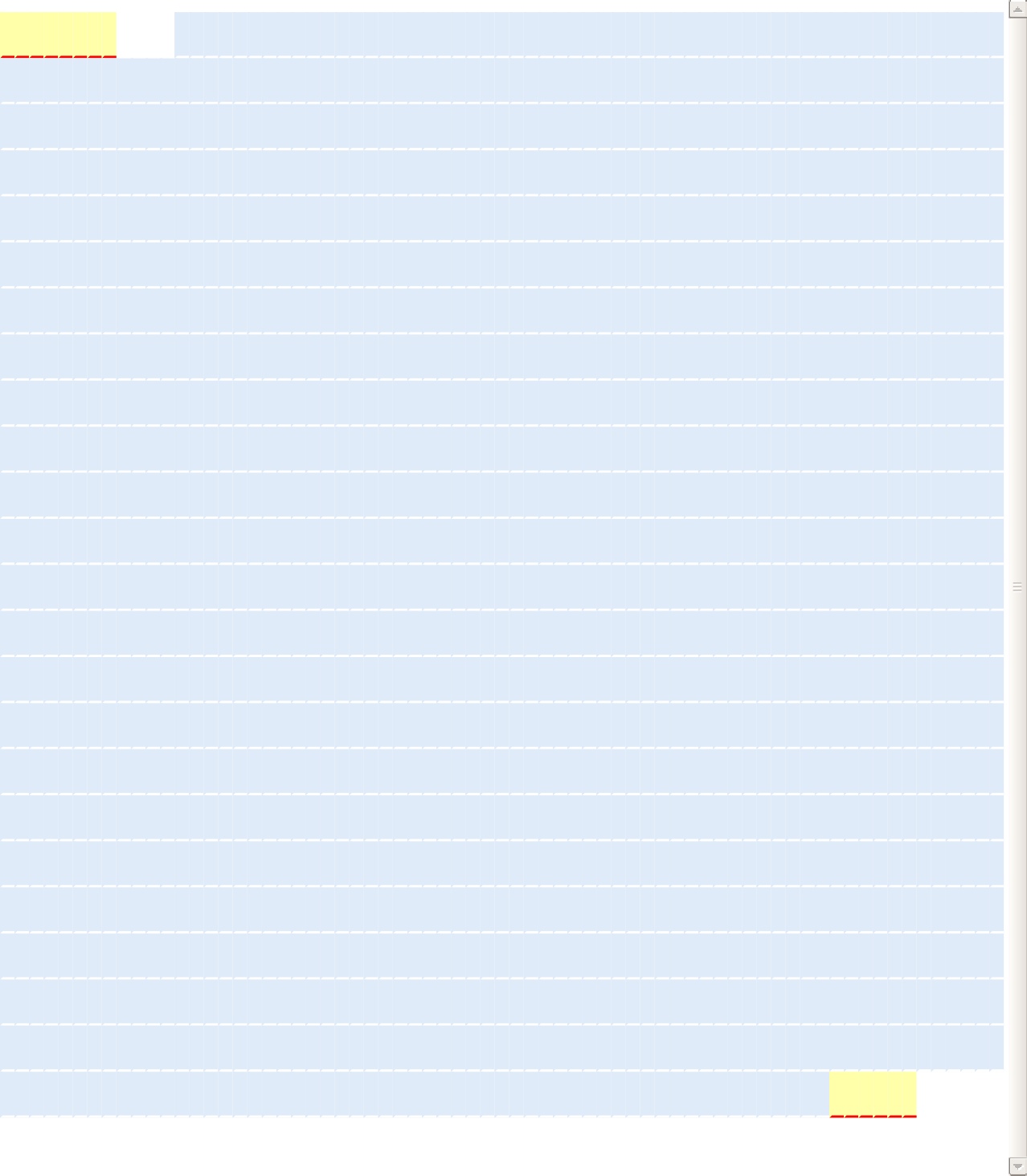
**M	A	S	R	K	E	N	A	K	S	A	N	R	V	L	R	I	S	Q**

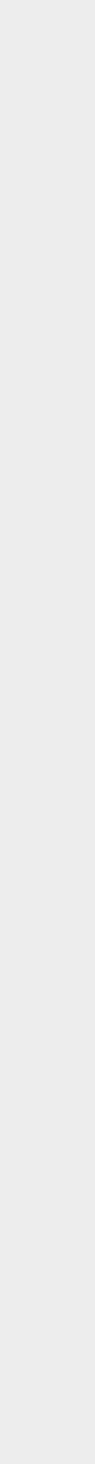

1.

70.

139.

208.

277.

346.

415.

484.

553.

622.

691.

760.

829.

898.

967.

1036.

1105.

1174.

1243.

1312.

1381.

1450.

1519.

1588.

G C G G C C G C C A A A A T G G C C A G C C G C A A G G A G A A T G C C A A G A G C G C C A A T C G C G T G C T G C G C A T C A G T C A G

**L	D	A**

**L	E	L	N	K	A	L**

**E	Q	L	V	W	S	Q	F	T	Q	C	F	H**

C T G G A T G C C C T G G A G C T G A A C A A G G C C C T G G A G C A G C T C G T G T G G T C C C A G T T C A C C C A G T G C T T C C A C

**G	F	K**

**P	G	L	L	A	R	F**

**E	P	E	V	K	A	C	L	W	V	F	L	W**

G G C T T C A A G C C A G G A C T G C T G G C C C G C T T T G A G C C C G A A G T G A A G G C C T G C C T G T G G G T G T T C C T G T G G

**R	F	T**

**I	Y	S	K	N	A	T**

**V	G	Q	S	V	L	N	I	K	Y	K	N	D**

C G C T T C A C C A T C T A C A G C A A G A A C G C C A C C G T G G G C C A G A G C G T G C T G A A C A T C A A G T A C A A G A A C G A C

**F	S	P**

**N	L	R	Y	Q	P	P**

**S	K	N	Q	K	I	W	Y	A	V	C	T**	**I**

T T C A G C C C C A A C C T G C G C T A C C A G C C C C C G A G C A A G A A C C A G A A G A T T T G G T A C G C C G T G T G C A C C A T C

**G	G	R**

**W	L	E	E	R	C	Y**

**D	L	F	R	N	H	H	L	A	S	F	G	K**

G G C G G A C G C T G G C T G G A G G A G C G C T G C T A C G A T C T G T T C C G C A A C C A C C A C C T G G C C A G C T T C G G C A A A

**V	K	Q**

**C	V	N	F	V	I	G**

**L	L	K	L	G	G	L	I	N	F	L	I	F**

G T G A A G C A G T G C G T G A A C T T C G T G A T C G G C C T G C T G A A G C T G G G C G G C C T G A T C A A C T T C C T G A T C T T C

**L	Q	R**

**G	K	F	A	T	L	T**

**E	R	L	L	G	I	H	S	V	F	C	K	P**

C T G C A G C G C G G C A A G T T C G C C A C C C T G A C C G A G C G C C T G C T G G G C A T T C A T A G C G T G T T C T G C A A G C C C

**Q	N**	**I**

**R	E	V	G	F	E	Y**

**M	N	R	E	L	L	W	H	G	F	A	E	F**

C A G A A C A T C C G C G A A G T G G G C T T C G A G T A C A T G A A C C G C G A G C T G C T G T G G C A C G G C T T C G C C G A G T T T

**L	I	F**

**L	L	P	L	I	N**	**V**

**Q	K	L	K	A	K	L	S	S	W	C	I	P**

C T G A T T T T C C T G C T G C C G C T G A T C A A C G T G C A G A A G C T G A A G G C C A A G C T G A G C A G C T G G T G C A T C C C A

**L	T	G**

**A	P	N	S	D	N	T**

**L	A	T	S	G	K	E	C	A	L	R	G	E**

C T G A C G G G A G C C C C C A A C A G C G A T A A C A C C C T G G C C A C C A G C G G A A A G G A G T G C G C C C T G C G C G G A G A G

**W	P	T**

**M	P	H	T	I	G	C**

**E	H	I	F	C	Y	F	C	A	K	S	S	F**

T G G C C A A C C A T G C C A C A C A C C A T T G G C T G C G A G C A C A T C T T C T G C T A C T T T T G C G C C A A G A G C A G C T T C

**L	F	D**

**V	Y	F	T	C	P	K**

**C	G	T	E	V	H	S	L	Q	P	L	K	S**

C T G T T C G A C G T G T A C T T C A C G T G C C C C A A G T G C G G C A C C G A G G T G C A C A G T C T G C A G C C A C T G A A G T C C

**G	I	E**

**M	S	E	V	N	A	L**

**V	S	K	G	E	E	L	F	T	G	V	V	P**

G G C A T C G A G A T G A G C G A A G T G A A C G C C C T G G T G T C C A A G G G C G A G G A G C T G T T T A C C G G C G T G G T G C C C

**I	L**	**V**

E **L	D	G	D	V	N**

**G	H	K	F	S	V	S	G	E	G	E	G	D**

A T T C T G G T G G A G C T G G A T G G C G A C G T G A A C G G C C A C A A G T T C A G C G T G T C C G G C G A G G G C G A G G G C G A C

**A	T	Y**

**G	K	L	T	L	K	F**

**I	C	T	T	G	K	L	P	V	P	W	P	T**

G C C A C C T A T G G A A A G C T G A C C C T G A A G T T C A T C T G C A C C A C C G G C A A G C T G C C C G T G C C A T G G C C A A C C

**L	V	T**

**T	L	T	Y	G	V	Q**

C **F	S	R	Y	P	D	H	M	K	Q	H	D**

C T C G T G A C C A C G C T G A C C T A T G G C G T G C A G T G C T T C A G C C G C T A C C C C G A T C A C A T G A A G C A G C A C G A T

F **F	K**

**S	A	M	P	E	G	Y**

**V	Q	E	R	T	I	F	F	K	D	D	G	N**

T T C T T C A A G T C C G C C A T G C C C G A G G G C T A C G T G C A G G A G C G C A C C A T C T T T T T C A A G G A T G A C G G C A A C

**Y	K	T**

**R	A	E	V	K	F	E**

G **D	T	L	V	N	R	I	E	L	K	G**	**I**

T A C A A G A C C C G C G C C G A A G T G A A G T T C G A G G G C G A T A C C C T C G T G A A C C G C A T C G A G C T G A A G G G C A T C

D **F	K**
E **D	G	N	I	L	G**
H **K	L	E	Y	N	Y	N	S	H	N	V	Y**

G A T T T C A A G G A G G A T G G A A A C A T C C T G G G C C A C A A G C T G G A G T A C A A C T A C A A C A G C C A C A A C G T G T A C

I **M	A**

**D	K	Q	K	N	G**	**I**

**K	V	N	F	K	I	R	H	N	I	E	D	G**

A T C A T G G C C G A C A A G C A G A A G A A C G G C A T C A A A G T G A A C T T C A A G A T T C G C C A C A A C A T C G A G G A T G G C

**S	V	Q**

**L	A	D	H	Y	Q	Q**

**N	T	P	I	G	D	G	P	V	L	L	P	D**

A G C G T G C A G C T G G C C G A C C A C T A C C A G C A G A A C A C C C C C A T C G G A G A T G G C C C C G T G C T G C T G C C C G A T

**N	H	Y**

**L	S	T	Q	S	A	L**

**S	K	D	P	N	E	K	R	D	H	M	V	L**

A A C C A C T A C C T G A G T A C C C A G A G C G C C C T G A G C A A G G A T C C C A A C G A G A A G C G C G A C C A C A T G G T G C T G

**L	E	F**

**V	T	A	A	G	I	T**

L **G	M	D	E	L	Y	K	***

C T G G A G T T T G T G A C C G C C G C C G G C A T T A C C C T G G G C A T G G A T G A G C T G T A C A A G T A G C T C G A G

The histograms show the percentage of sequence codons which fall into a certain quality class. The quality value of the most frequently used codon for a given amino acid in the desired expression system is set to 100, the remaining codons are scaled accordingly (see also Sharp, P.M., Li, W.H., Nucleic Acids Res. 15 (3),1987).

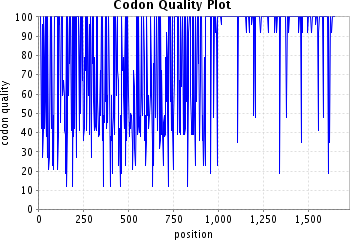

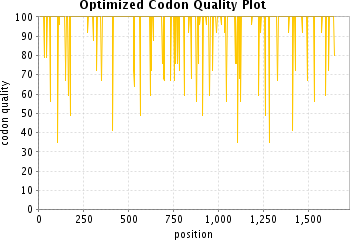

The plots show the quality of the used codon at the indicated codon position.

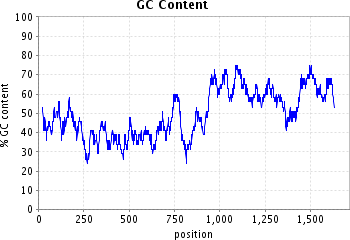

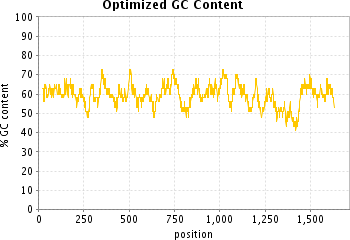

The plots show the GC content in a 40 bp window centered at the indicated nucleotide position.

**Sequence name / optimized for**

#### PEX16_human/ Drosophila melanogaster

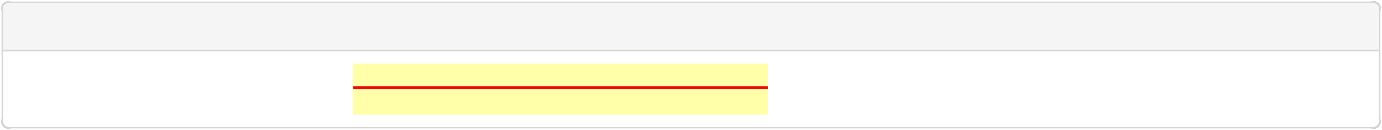

**ORF**

**Protected sites**

**Protected areas**

**Motifs to avoid**

13-1065 [ATG...TAG]

NotI [GCGGCCGC] XhoI [CTCGAG]

1-8 NotI [GCGGCCGC]

1066-1071 XhoI [CTCGAG]

M
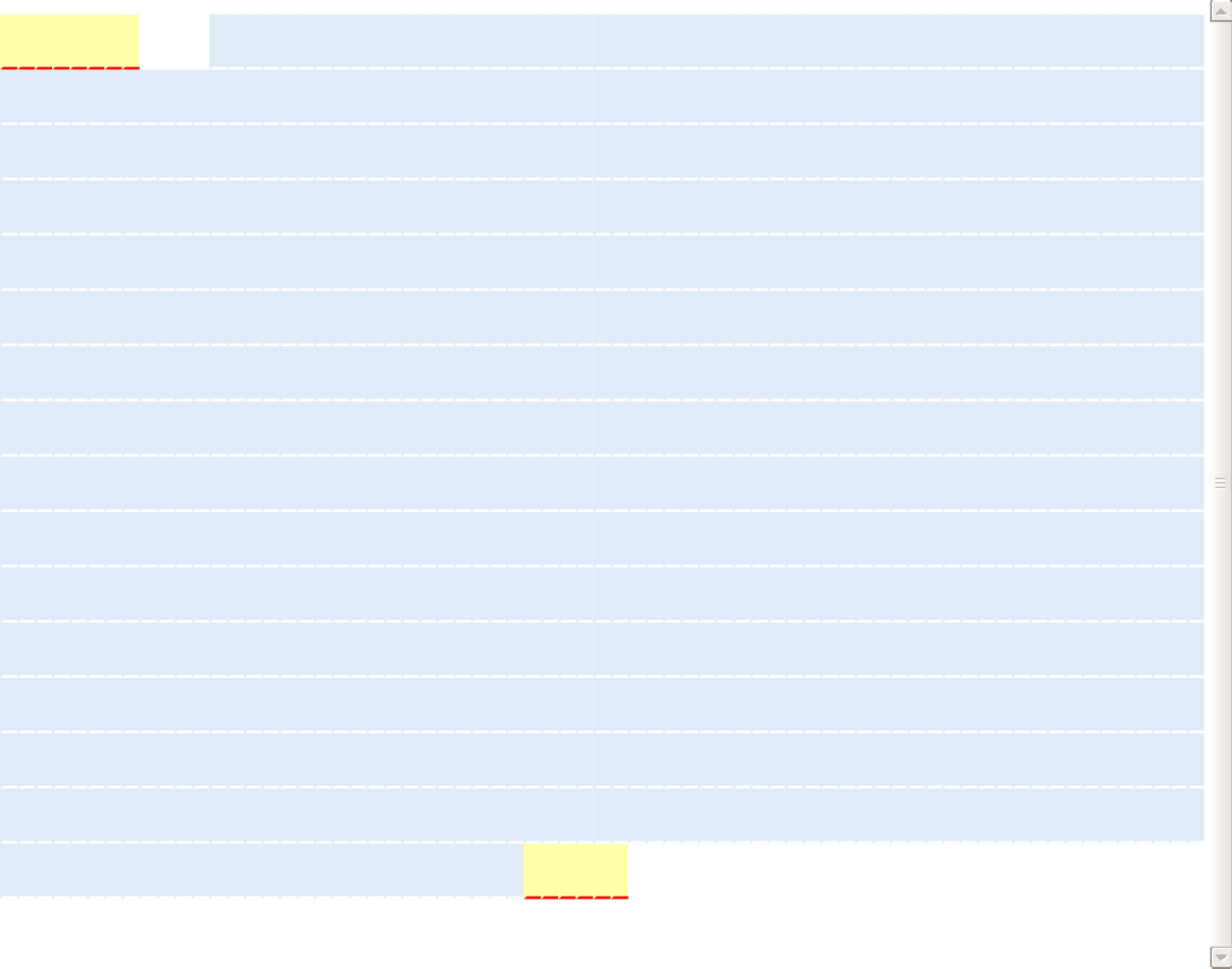
**G	K	P	I	P	N	P	L	L	G	L	D	S	T	E	K	L	R**

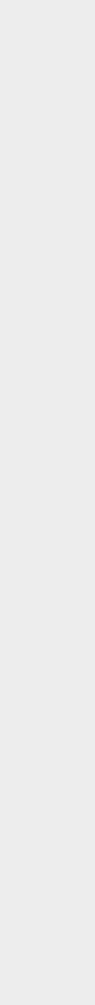

1.

70.

139.

208.

277.

346.

415.

484.

553.

622.

691.

760.

829.

898.

967.

1036.

G C G G C C G C C A A A A T G G G C A A G C C C A T C C C C A A T C C A C T G C T G G G C C T G G A T A G C A C C G A G A A G C T G C G C

**L	L	G**

**L	R	Y	Q	E	Y**	**V**

**T	R	H	P	A	A	T	A	Q	L	E	T	A**

C T G C T G G G A C T G C G C T A C C A G G A G T A T G T G A C C C G C C A T C C A G C C G C C A C C G C C C A G C T G G A G A C C G C C

**V	R	G**

F **S	Y	L	L	A	G**
R **F	A	D	S	H	E	L	S	E	L	V	Y**

G T G C G C G G A T T T T C C T A T C T G C T G G C C G G A C G C T T C G C C G A T A G C C A C G A G C T G A G C G A G C T G G T G T A C

S **A	S**

**N	L	L	V	L	L	N**

**D	G	I	L	R	K	E	L	R	K	K	L	P**

A G C G C C A G C A A C C T G C T G G T G C T G C T G A A C G A T G G C A T C C T G C G C A A G G A G C T G C G C A A G A A G C T G C C C

**V	S	L**

S **Q	Q	K	L	L	T**

**W	L	S	V	L	E	C	V	E	V	F	M	E**

G T G T C C C T G A G C C A G C A G A A G C T G C T G A C C T G G C T G T C C G T G C T G G A G T G C G T G G A G G T G T T C A T G G A G

**M	G	A**

**A	K	V	W	G	E**	**V**

G **R	W	L	V	I	A	L	I	Q	L	A	K**

A T G G G A G C C G C C A A A G T G T G G G G C G A A G T G G G A C G C T G G C T C G T G A T C G C C C T G A T C C A G C T G G C C A A G

**A	V	L**

**R	M	L	L	L	L	W**

**F	K	A	G	L	Q	T	S	P	P	I	V	P**

G C C G T G C T G C G C A T G T T G C T G C T G C T G T G G T T C A A G G C C G G A C T G C A G A C C A G C C C C C C A A T C G T G C C A

**L	D	R**

**E	T	Q	A	Q	P	P**

**D	G	D	H	S	P	G	N	H	E	Q	S	Y**

C T G G A T C G C G A G A C G C A G G C C C A G C C A C C A G A T G G C G A T C A C T C C C C A G G C A A C C A C G A G C A G A G C T A C

**V	G	K**

**R	S	N	R	V	V	R**

**T	L	Q	N	T	P	S	L	H	S	R	H	W**

G T G G G C A A G C G C T C C A A T C G C G T C G T G C G C A C C C T G C A G A A T A C C C C C A G T C T G C A C A G C C G C C A T T G G

**G	A	P**

**Q	Q	R	E	G	R	Q**

**Q	Q	H	H	E	E	L	S	A	T	P	T	P**

G G A G C C C C C C A G C A G C G C G A G G G A C G C C A G C A G C A G C A T C A C G A G G A G C T G A G T G C C A C C C C A A C C C C A

**L	G	L**

**Q	E	T	I	A	E	F**

**L	Y	I	A	R	P	L	L	H	L	L	S	L**

C T G G G A C T G C A G G A G A C G A T C G C C G A G T T C C T G T A C A T T G C C C G C C C A C T G C T G C A T C T G C T G A G C C T G

**G	L	W**

**G	Q	R	S	W	K	P**

**W	L	L	A	G	V	V	D	V	T	S	L	S**

G G A C T G T G G G G A C A G C G C A G T T G G A A G C C C T G G C T G C T G G C C G G C G T G G T G G A T G T G A C C A G T C T G A G C

**L	L	S**

**D	R	K	G	L	T	R**

**R	E	R	R	E	L	R	R	R	T	I	L	L**

C T G C T G A G C G A T C G C A A G G G A C T G A C C C G C C G C G A G C G C C G C G A G C T G C G C C G C C G C A C C A T T C T G C T G

**L	Y	Y**

**L	L	R	S	P	F	Y**

**D	R	F	S	E	A	R	I	L	F	L	L	Q**

C T G T A C T A T C T G C T G C G C T C C C C C T T C T A C G A T C G C T T C T C C G A G G C C C G C A T C C T G T T T C T G C T G C A G

**L	L	A**

**D	H	V	P	G	V	G**

L **V	T	R	P	L	M	D	Y	L	P	T	W**

C T G C T G G C C G A T C A C G T G C C C G G C G T G G G A C T C G T G A C G C G C C C A C T G A T G G A T T A C C T G C C C A C C T G G

**Q	K	I	Y	F	Y	S	W	G	***

C A G A A G A T C T A C T T C T A C A G C T G G G G C T A G C T C G A G

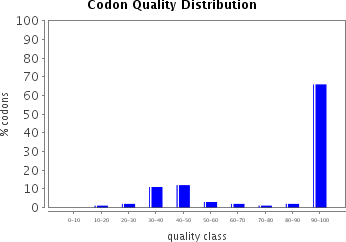

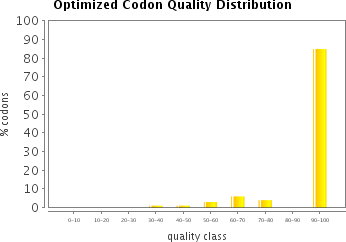

The histograms show the percentage of sequence codons which fall into a certain quality class. The quality value of the most frequently used codon for a given amino acid in the desired expression system is set to 100, the remaining codons are scaled accordingly (see also Sharp, P.M., Li, W.H., Nucleic Acids Res. 15 (3),1987).

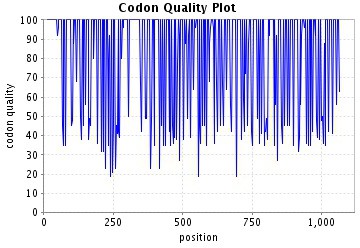

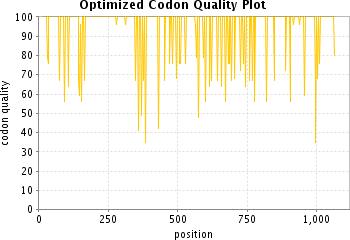

The plots show the quality of the used codon at the indicated codon position.

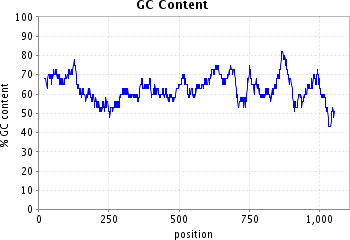

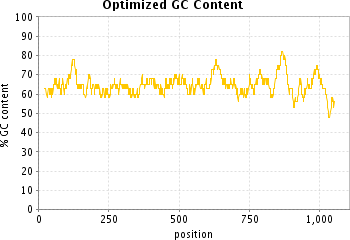

The plots show the GC content in a 40 bp window centered at the indicated nucleotide position.

**Sequence name / optimized for**

#### PEX16_R176X/ Drosophila melanogaster

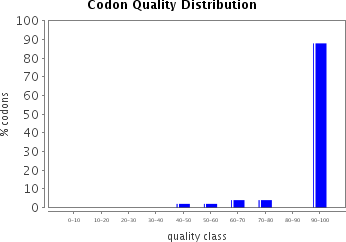

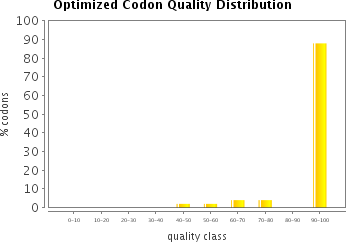

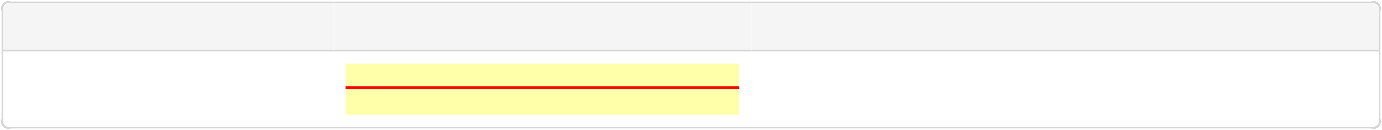

**ORF**

**Protected sites**

**Protected areas**

**Motifs to avoid**

13-582 [ATG...TAA]

NotI [GCGGCCGC] XhoI [CTCGAG]

1-8 NotI [GCGGCCGC]

583-588 XhoI [CTCGAG]

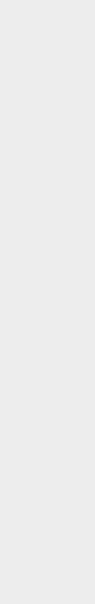

1.

70.

139.

208.

277.

346.

415.

484.

553.

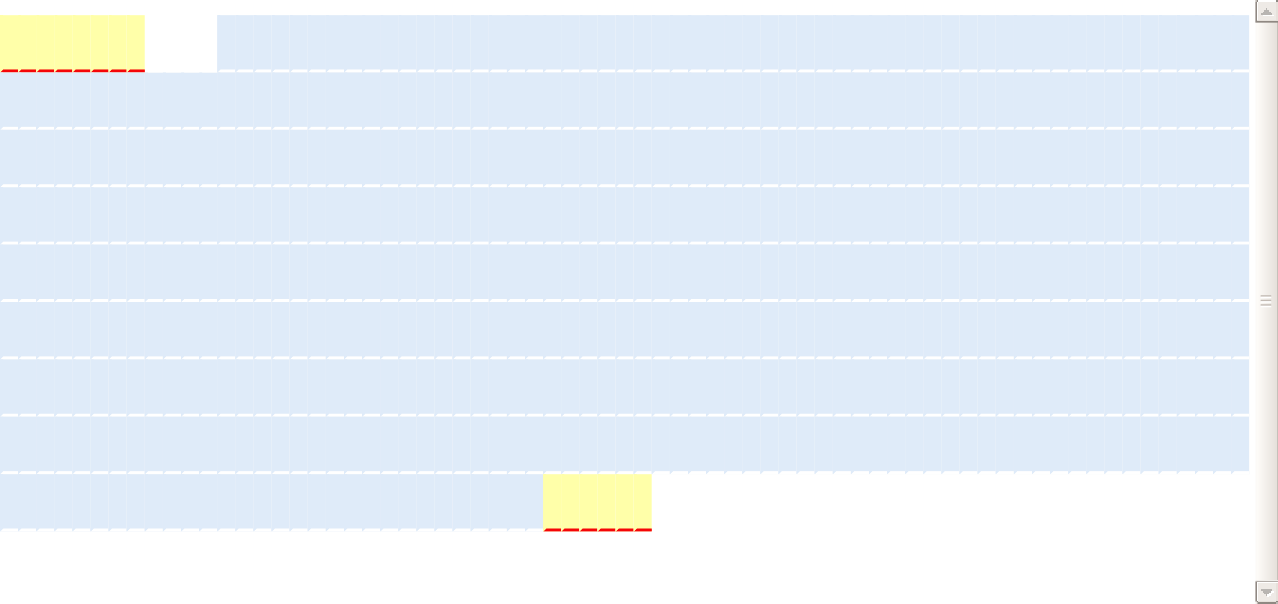

**M	G	K	P	I	P	N	P	L	L	G	L	D	S	T	E	K	L	R**

G C G G C C G C C A A A A T G G G C A A G C C C A T C C C C A A T C C A C T G C T G G G C C T G G A T A G C A C C G A G A A G C T G C G C

**L	L	G	L	R	Y	Q	E	Y	V	T	R	H	P	A	A	T	A	Q	L	E	T	A**

C T G C T G G G A C T G C G C T A C C A G G A G T A T G T G A C C C G C C A T C C A G C C G C C A C C G C C C A G C T G G A G A C C G C C

**V	R	G	F	S	Y	L	L	A	G	R	F	A	D	S	H	E	L	S	E	L	V	Y**

G T G C G C G G A T T T T C C T A T C T G C T G G C C G G A C G C T T C G C C G A T A G C C A C G A G C T G A G C G A G C T G G T G T A C

**S	A	S	N	L	L	V	L	L	N	D	G	I	L	R	K	E	L	R	K	K	L	P**

A G C G C C A G C A A C C T G C T G G T G C T G C T G A A C G A T G G C A T C C T G C G C A A G G A G C T G C G C A A G A A G C T G C C C

**V	S	L	S	Q	Q	K	L	L	T	W	L	S	V	L	E	C	V	E	V	F	M	E**

G T G T C C C T G A G C C A G C A G A A G C T G C T G A C C T G G C T G T C C G T G C T G G A G T G C G T G G A G G T G T T C A T G G A G

**M	G	A	A	K	V	W	G	E	V	G	R	W	L	V	I	A	L	I	Q	L	A	K**

A T G G G A G C C G C C A A A G T G T G G G G C G A A G T G G G A C G C T G G C T C G T G A T C G C C C T G A T C C A G C T G G C C A A G

**A	V	L	R	M	L	L	L	L	W	F	K	A	G	L	Q	T	S	P	P	I	V	P**

G C C G T G C T G C G C A T G T T G C T G C T G C T G T G G T T C A A G G C C G G A C T G C A G A C C A G C C C C C C A A T C G T G C C A

**L	D	R	E	T	Q	A	Q	P	P	D	G	D	H	S	P	G	N	H	E	Q	S	Y**

C T G G A T C G C G A G A C G C A G G C C C A G C C A C C A G A T G G C G A T C A C T C C C C A G G C A A C C A C G A G C A G A G C T A C

**V	G	K	R	S	N	R	V	V	***

G T G G G C A A G C G C T C C A A C C G C G T G G T G T A A C T C G A G

The histograms show the percentage of sequence codons which fall into a certain quality class. The quality value of the most frequently used codon for a given amino acid in the desired expression system is set to 100, the remaining codons are scaled accordingly (see also Sharp, P.M., Li, W.H., Nucleic Acids Res. 15 (3),1987).

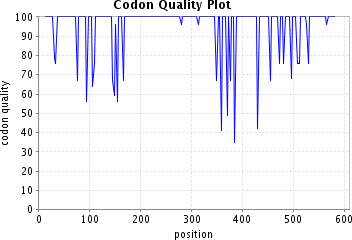

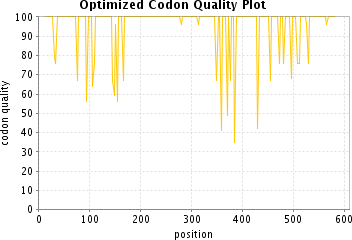

The plots show the quality of the used codon at the indicated codon position.

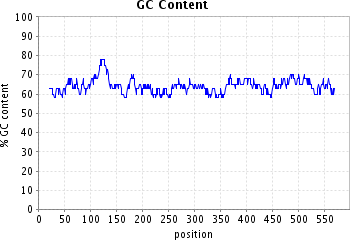

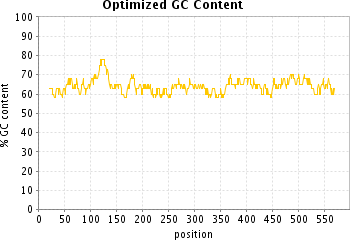

The plots show the GC content in a 40 bp window centered at the indicated nucleotide position.

**Sequence name / optimized for**

#### PEX16_del_955TCT/ Drosophila melanogaster

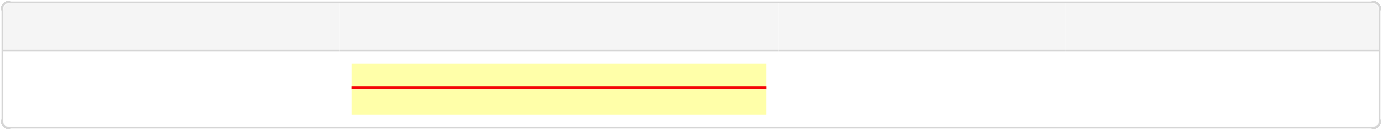

**ORF**

**Protected sites**

**Protected areas**

**Motifs to avoid**

13-1062 [ATG...TAA]

NotI [GCGGCCGC] XhoI [CTCGAG]

1-8 NotI [GCGGCCGC]

1063-1068 XhoI [CTCGAG]

M
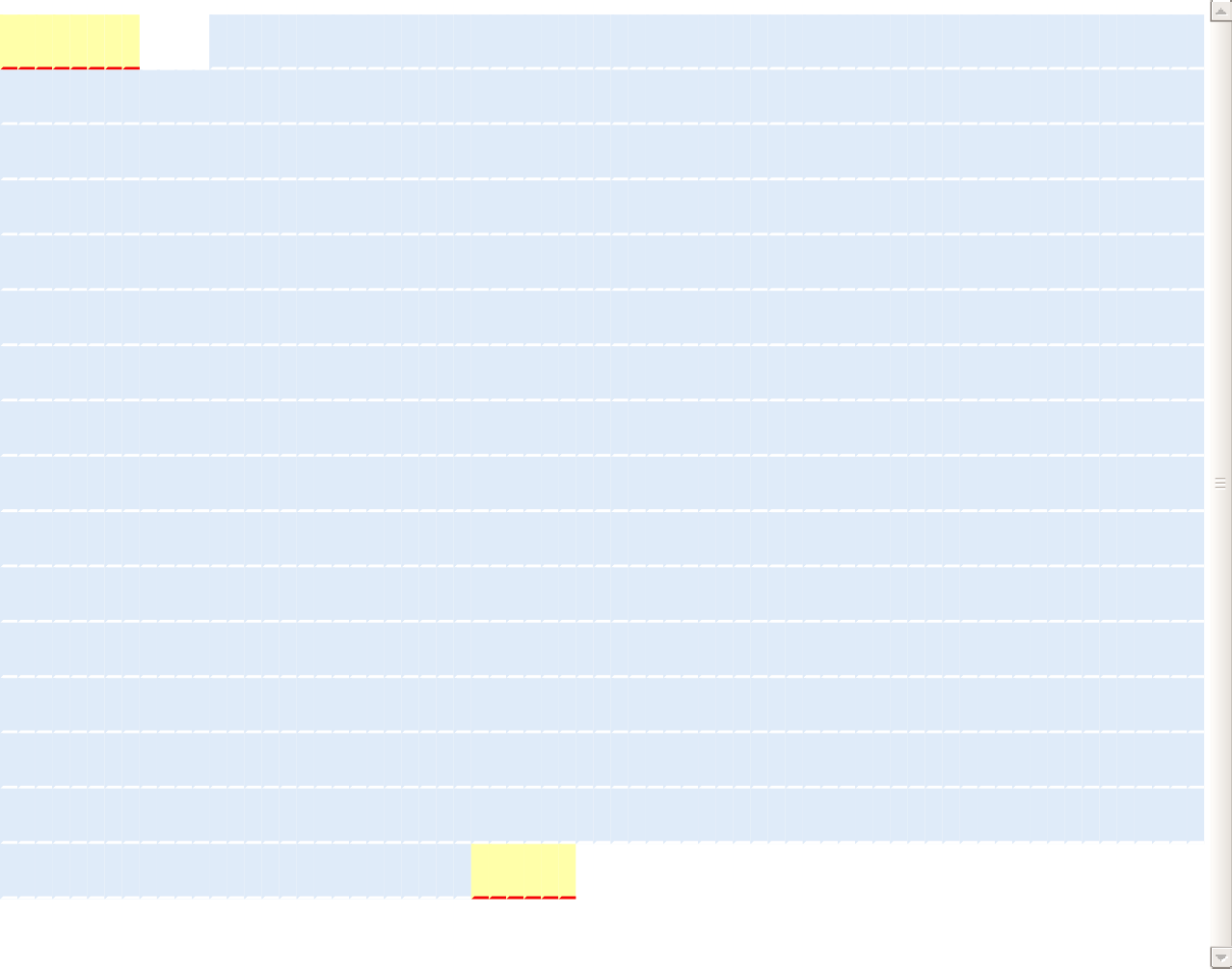
**G	K	P	I	P	N	P	L	L	G	L	D	S	T	E	K	L	R**

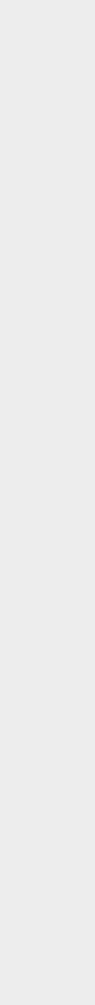

1.

70.

139.

208.

277.

346.

415.

484.

553.

622.

691.

760.

829.

898.

967.

1036.

G C G G C C G C C A A A A T G G G C A A G C C C A T C C C C A A T C C A C T G C T G G G C C T G G A T A G C A C C G A G A A G C T G C G C

**L	L	G**

**L	R	Y	Q	E	Y**	**V**

T **R	H	P	A	A	T	A	Q	L	E	T	A**

C T G C T G G G A C T G C G C T A C C A G G A G T A T G T G A C C C G C C A T C C A G C C G C C A C C G C C C A G C T G G A G A C C G C C

**V	R	G**

**F	S	Y	L	L	A	G**

**R	F	A	D	S	H	E	L	S	E	L	V	Y**

G T G C G C G G A T T T T C C T A T C T G C T G G C C G G A C G C T T C G C C G A T A G C C A C G A G C T G A G C G A G C T G G T G T A C

**S	A	S**

**N	L	L	V	L	L	N**

**D	G	I	L	R	K	E	L	R	K	K	L	P**

A G C G C C A G C A A C C T G C T G G T G C T G C T G A A C G A T G G C A T C C T G C G C A A G G A G C T G C G C A A G A A G C T G C C C

**V	S	L**

**S	Q	Q	K	L	L	T**

**W	L	S	V	L	E	C	V	E	V	F	M	E**

G T G T C C C T G A G C C A G C A G A A G C T G C T G A C C T G G C T G T C C G T G C T G G A G T G C G T G G A G G T G T T C A T G G A G

**M	G	A**

**A	K	V	W	G	E**	**V**

**G	R	W	L	V	I	A	L	I	Q	L	A	K**

A T G G G A G C C G C C A A A G T G T G G G G C G A A G T G G G A C G C T G G C T C G T G A T C G C C C T G A T C C A G C T G G C C A A G

**A	V	L**

**R	M	L	L	L	L	W**

**F	K	A	G	L	Q	T	S	P	P	I	V	P**

G C C G T G C T G C G C A T G T T G C T G C T G C T G T G G T T C A A G G C C G G A C T G C A G A C C A G C C C C C C A A T C G T G C C A

**L	D	R**

**E	T	Q	A	Q	P	P**

**D	G	D	H	S	P	G	N	H	E	Q	S	Y**

C T G G A T C G C G A G A C G C A G G C C C A G C C A C C A G A T G G C G A T C A C T C C C C A G G C A A C C A C G A G C A G A G C T A C

**V	G	K**

**R	S	N	R	V	V	R**

**T	L	Q	N	T	P	S	L	H	S	R	H	W**

G T G G G C A A G C G C T C C A A T C G C G T C G T G C G C A C C C T G C A G A A T A C C C C C A G T C T G C A C A G C C G C C A T T G G

**G	A	P**

**Q	Q	R	E	G	R	Q**

**Q	Q	H	H	E	E	L	S	A	T	P	T	P**

G G A G C C C C C C A G C A G C G C G A G G G A C G C C A G C A G C A G C A T C A C G A G G A G C T G A G T G C C A C C C C A A C C C C A

**L	G	L**

**Q	E	T	I	A	E	F**

**L	Y	I	A	R	P	L	L	H	L	L	S	L**

C T G G G A C T G C A G G A G A C G A T C G C C G A G T T C C T G T A C A T T G C C C G C C C A C T G C T G C A T C T G C T G A G C C T G

**G	L	W**

**G	Q	R	S	W	K	P**

**W	L	L	A	G	V	V	D	V	T	S	L	S**

G G A C T G T G G G G A C A G C G C A G T T G G A A G C C C T G G C T G C T G G C C G G C G T G G T G G A T G T G A C C A G T C T G A G C

**L	L	S**

**D	R	K	G	L	T	R**

**R	E	R	R	E	L	R	R	R	T	I	L	L**

C T G C T G A G C G A T C G C A A G G G A C T G A C C C G C C G C G A G C G C C G C G A G C T G C G C C G C C G C A C C A T T C T G C T G

**L	Y	Y**

**L	L	R	S	P	F	Y**

**D	R	F	S	E	A	R	I	L	F	L	L	Q**

C T G T A C T A T C T G C T G C G C T C C C C C T T C T A C G A T C G C T T C T C C G A G G C C C G C A T C C T G T T T C T G C T G C A G

**L	L	A**

**D	H	V	P	G	V	G**

**L	V	T	R	P	L	M	D	Y	L	P	T	W**

C T G C T G G C C G A T C A C G T G C C C G G C G T G G G A C T C G T G A C G C G C C C A C T G A T G G A T T A C C T G C C C A C C T G G

**Q	K	I	Y	Y	S	W	G	***

C A G A A G A T C T A C T A C A G C T G G G G C T A A C T C G A G

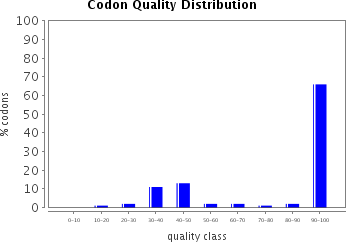

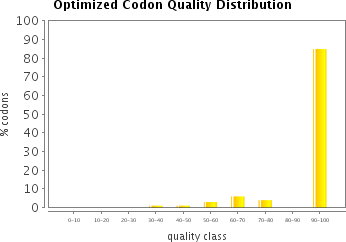

The histograms show the percentage of sequence codons which fall into a certain quality class. The quality value of the most frequently used codon for a given amino acid in the desired expression system is set to 100, the remaining codons are scaled accordingly (see also Sharp, P.M., Li, W.H., Nucleic Acids Res. 15 (3),1987).

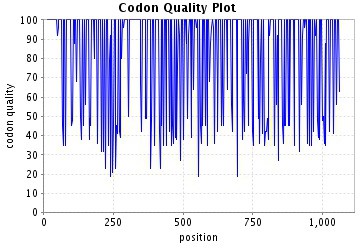

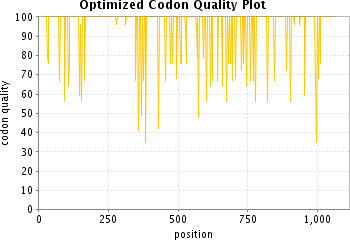

The plots show the quality of the used codon at the indicated codon position.

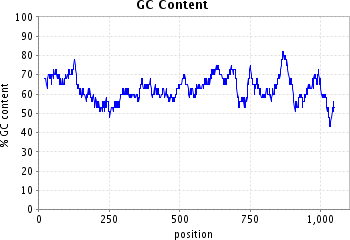

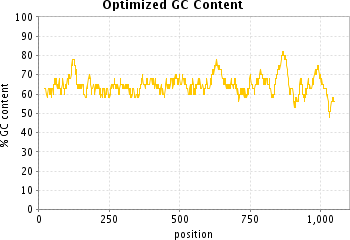

The plots show the GC content in a 40 bp window centered at the indicated nucleotide position.

MASRKENAKSANRVLRISQLDALELNKALEQLVWSQFTQCFHGFKPGLLARFEPEV 56

KACLWVFLWRFTIYSKNATVGQSVLNIKYKNDFSPNLRYQPPSKNQKIWYAVCTIGGRWL 116

EERCYDLFRNHHLASFGKVKQCVNFVIGLLKLGGLINFLIFLQRGKFATLTERLLGIHSV 176

FCKPQNIREVGFEYMNRELLWHGFAEFLIFLLPLINVQKLKAKLSSWCIPLTGAPNSDNT 236

LATSGKECALCGEWPTMPHTIGCEHIFCYFCAKSSFLFDVYFTCPKCGTEVHSLQPLKSG 296

IEMSEVNALVSKGEELFTGVVPILVELDGDVNGHKFSVSGEGEGDATYGKLTLKFICTTG 356

KLPVPWPTLVTTLTYGVQCFSRYPDHMKQHDFFKSAMPEGYVQERTIFFKDDGNYKTRAE 416

VKFEGDTLVNRIELKGIDFKEDGNILGHKLEYNYNSHNVYIMADKQKNGIKVNFKIRHNI 476

EDGSVQLADHYQQNTPIGDGPVLLPDNHYLSTQSALSKDPNEKRDHMVLLEFVTAAGITL 536

GMDELYK 543

### PEX2_E55K

MASRKENAKSANRVLRISQLDALELNKALEQLVWSQFTQCFHGFKPGLLARFEPKV 56

KACLWVFLWRFTIYSKNATVGQSVLNIKYKNDFSPNLRYQPPSKNQKIWYAVCTIGGRWL 116

EERCYDLFRNHHLASFGKVKQCVNFVIGLLKLGGLINFLIFLQRGKFATLTERLLGIHSV 176

FCKPQNIREVGFEYMNRELLWHGFAEFLIFLLPLINVQKLKAKLSSWCIPLTGAPNSDNT 236

LATSGKECALCGEWPTMPHTIGCEHIFCYFCAKSSFLFDVYFTCPKCGTEVHSLQPLKSG 296

IEMSEVNALVSKGEELFTGVVPILVELDGDVNGHKFSVSGEGEGDATYGKLTLKFICTTG 356

KLPVPWPTLVTTLTYGVQCFSRYPDHMKQHDFFKSAMPEGYVQERTIFFKDDGNYKTRAE 416

VKFEGDTLVNRIELKGIDFKEDGNILGHKLEYNYNSHNVYIMADKQKNGIKVNFKIRHNI 476

EDGSVQLADHYQQNTPIGDGPVLLPDNHYLSTQSALSKDPNEKRDHMVLLEFVTAAGITL 536

GMDELYK 543

### PEX2_C247R

MASRKENAKSANRVLRISQLDALELNKALEQLVWSQFTQCFHGFKPGLLARFEPEV 56

KACLWVFLWRFTIYSKNATVGQSVLNIKYKNDFSPNLRYQPPSKNQKIWYAVCTIGGRWL 116

EERCYDLFRNHHLASFGKVKQCVNFVIGLLKLGGLINFLIFLQRGKFATLTERLLGIHSV 176

FCKPQNIREVGFEYMNRELLWHGFAEFLIFLLPLINVQKLKAKLSSWCIPLTGAPNSDNT 236

LATSGKECALRGEWPTMPHTIGCEHIFCYFCAKSSFLFDVYFTCPKCGTEVHSLQPLKSG 296

IEMSEVNALVSKGEELFTGVVPILVELDGDVNGHKFSVSGEGEGDATYGKLTLKFICTTG 356

KLPVPWPTLVTTLTYGVQCFSRYPDHMKQHDFFKSAMPEGYVQERTIFFKDDGNYKTRAE 416

VKFEGDTLVNRIELKGIDFKEDGNILGHKLEYNYNSHNVYIMADKQKNGIKVNFKIRHNI 476

EDGSVQLADHYQQNTPIGDGPVLLPDNHYLSTQSALSKDPNEKRDHMVLLEFVTAAGITL 536

GMDELYK 543

### PEX2_W223X

MASRKENAKSANRVLRISQLDALELNKALEQLVWSQFTQCFHGFKPGLLARFEPEV 56

KACLWVFLWRFTIYSKNATVGQSVLNIKYKNDFSPNLRYQPPSKNQKIWYAVCTIGGRWL 116

EERCYDLFRNHHLASFGKVKQCVNFVIGLLKLGGLINFLIFLQRGKFATLTERLLGIHSV 176

FCKPQNIREVGFEYMNRELLWHGFAEFLIFLLPLINVQKLKAKLSSVSKGEELFTGVVPI 236

LVELDGDVNGHKFSVSGEGEGDATYGKLTLKFICTTGKLPVPWPTLVTTLTYGVQCFSRY 296

PDHMKQHDFFKSAMPEGYVQERTIFFKDDGNYKTRAEVKFEGDTLVNRIELKGIDFKEDG 356

NILGHKLEYNYNSHNVYIMADKQKNGIKVNFKIRHNIEDGSVQLADHYQQNTPIGDGPVL 416

LPDNHYLSTQSALSKDPNEKRDHMVLLEFVTAAGITLGMDELYK 460

### PEX2_R119X

MASRKENAKSANRVLRISQLDALELNKALEQLVWSQFTQCFHGFKPGLLARFEPEV 56

KACLWVFLWRFTIYSKNATVGQSVLNIKYKNDFSPNLRYQPPSKNQKIWYAVCTIGGRWL 116

EEVSKGEELFTGVVPILVELDGDVNGHKFSVSGEGEGDATYGKLTLKFICTTGKLPVPWP 176

TLVTTLTYGVQCFSRYPDHMKQHDFFKSAMPEGYVQERTIFFKDDGNYKTRAEVKFEGDT 236

LVNRIELKGIDFKEDGNILGHKLEYNYNSHNVYIMADKQKNGIKVNFKIRHNIEDGSVQL 296

ADHYQQNTPIGDGPVLLPDNHYLSTQSALSKDPNEKRDHMVLLEFVTAAGITLGMDELYK 356

MGKPIPNPLLGLDSTEKLRLLGLRYQEYVTRHPAATAQLETAVRGFSYLLAGRFAD 56

SHELSELVYSASNLLVLLNDGILRKELRKKLPVSLSQQKLLTWLSVLECVEVFMEMGAAK 116

VWGEVGRWLVIALIQLAKAVLRMLLLLWFKAGLQTSPPIVPLDRETQAQPPDGDHSPGNH 176

EQSYVGKRSNRVVRTLQNTPSLHSRHWGAPQQREGRQQQHHEELSATPTPLGLQETIAEF 236

LYIARPLLHLLSLGLWGQRSWKPWLLAGVVDVTSLSLLSDRKGLTRRERRELRRRTILLL 296

YYLLRSPFYDRFSEARILFLLQLLADHVPGVGLVTRPLMDYLPTWQKIYFYSWG 350

PEX16_R176X

MGKPIPNPLLGLDSTEKLRLLGLRYQEYVTRHPAATAQLETAVRGFSYLLAGRFAD 56

SHELSELVYSASNLLVLLNDGILRKELRKKLPVSLSQQKLLTWLSVLECVEVFMEMGAAK 116

VWGEVGRWLVIALIQLAKAVLRMLLLLWFKAGLQTSPPIVPLDRETQAQPPDGDHSPGNH 176

EQSYVGKRSNRVV* 193

PEX16_del_955TCT

MGKPIPNPLLGLDSTEKLRLLGLRYQEYVTRHPAATAQLETAVRGFSYLLAGRFAD 56

SHELSELVYSASNLLVLLNDGILRKELRKKLPVSLSQQKLLTWLSVLECVEVFMEMGAAK 116

VWGEVGRWLVIALIQLAKAVLRMLLLLWFKAGLQTSPPIVPLDRETQAQPPDGDHSPGNH 176

EQSYVGKRSNRVVRTLQNTPSLHSRHWGAPQQREGRQQQHHEELSATPTPLGLQETIAEF 236

LYIARPLLHLLSLGLWGQRSWKPWLLAGVVDVTSLSLLSDRKGLTRRERRELRRRTILLL 296

YYLLRSPFYDRFSEARILFLLQLLADHVPGVGLVTRPLMDYLPTWQKIY_YSWG 349
